# Variations in the root-soil system influence the grapevine holobiont by shaping plant physiology and root microbiome

**DOI:** 10.1101/2025.08.14.670050

**Authors:** Massimo Guazzini, Ramona Marasco, Slobodanka Radović, Elisa Pellegrini, Marco Vuerich, Arianna Lodovici, Giorgia Dubsky De Wittenau, Eleonora Paparelli, Gabriele Magris, Laura Zanin, Marco Contin, Elisa De Luca, Daniele Daffonchio, Gabriele Di Gaspero, Fabio Marroni

## Abstract

Soil-dwelling bacteria and fungi play a crucial role in plant health and productivity by engaging in complex interactions that shape and are shaped by soil physico-chemical properties. In this study, we employed a multi-omics approach to investigate how variations in soil composition affect the grapevine holobiont. Grape plantlets were grown in three distinct soil types, namely sand, peat, and peat-manure. To further assess how variation in soil and root conditions affects the holobiont’s response, we included treatments involving soil autoclaving and root surface sterilisation across all soil types. We found that soil type significantly influences leaf multielement composition and concentration, while also shaping the bacterial and fungal communities associated with the plant rhizosphere. This shift led to changes in taxa involved in nitrogen fixation, biocontrol, and pathogenicity. Autoclaving soils consistently reduced bacterial diversity across all soil types, whereas fungal communities were less affected. In contrast, thermal treatment of roots had only a minor impact on microbial community composition but did induce transcriptional changes in the root and altered leaf macronutrient concentrations. Our findings indicate that differences in soil composition reshape the entire root-soil continuum, ultimately affecting plant physiology at multiple levels—from root function to leaf nutrient status. This highlights that the soil is not a passive growth medium but a key determinant of grape holobiont structure and function. These results reinforce the view that plant health and adaptation arise from integrated, dynamic interactions among the host, its associated microbiome, and the surrounding soil matrix.

## 1. Introduction

Soil bacteria and fungi establish close associations with plants and can significantly influence their health and productivity (Turner, James and Poole, 2013; Bettenfeld *et al*., 2022; Berg *et al*., 2024). This intimate relationship has led to the concept of the “plant holobiont”, which refers to the integrated biological entity formed by the plant host and its associated microbiota, functioning as a single ecological unit (Vandenkoornhuyse *et al*., 2015; Hassani *et al.,* 2018; Mesny *et al.,* 2023; Berg *et al*., 2024).

The soil microbiome is primarily shaped by soil physico-chemical properties (Islam *et al*., 2020), which also directly affect plant physiology by regulating nutrient availability, water retention, and structural support (Lanyon *et al.,* 2004; El-Ramady *et al*., 2014). However, plants themself are key determinant in shaping their microbial associations, particularly in the rhizosphere—the narrow zone of soil surrounding plant roots—and adjacent soil still influenced by roots (Berendsen *et al.,* 2012; Sánchez-Cañizares *et al*., 2017). Within this zone, plants actively recruit and select edaphic microorganisms through the release of root exudates and volatile organic compounds (Pascale *et al*., 2020). The composition of these exudates is also influenced by multiple environmental and climatic factors and varies among plant species and cultivars, thereby shaping the root microbiome and influencing plant-microbe interactions (Herz *et al*., 2018; Ghatak *et al*., 2022).

The interplay between plants and soil microbiota and its consequences for plant physiology, development, and resilience is an active and expanding field of research, particularly in grapevine (*Vitis vinifera*), a high-value crop cultivated worldwide (Sánchez-Cañizares *et al*., 2017; Bettenfeld *et al*., 2022). Grapevine is grown across a wide range of soil types that differ in physical and chemical properties, as well as in microbial load and composition (Marasco *et al*., 2013; Uyan *et al.,* 2023). Vines are typically propagated as rooted cuttings in nurseries, where they establish early symbiotic associations with endo- and exo-symbionts, which can result in either mutualistic or pathogenic relationships (Gramaje *et al*., 2022; Lade *et al*., 2022). To mitigate the risk of pathogenic associations, nurseries often apply treatments aimed at reducing disease incidence, such as root thermal treatment (Waite and May, 2005; Lade *et al*., 2022). However, these treatments can have a long-term impact on the structuring of the plant microbiome and physiology (Vukicevich *et al*., 2018). Once mature, plantlets are transplanted into vineyards where they encounter different environmental conditions and continue to develop complex and long-term interactions with the microorganisms living in the surrounding soil (Deloire *et al*., 2005; van Leeuwen, 2010).

The relationships between grapevine, soil, and their associated microbiomes are highly interconnected and interdependent. Numerous studies suggest that vineyard soil serves as the primary source of grapevine-associated microbiomes (Zarraonaindia *et al*., 2015). While previous studies have investigated how plant cultivars and soil types influence the composition of the grapevine microbiome (Rolli *et al*., 2017; Nerva *et al*., 2021; Marasco *et al*., 2022), these approaches often overlook the systemic feedbacks that define holobiont response, i.e., plant and microbiome. Recognizing this, we designed a multi-omics framework that simultaneously interrogates soil physico-chemical properties, microbial community structure (both bacterial and fungal), and grapevine physiological responses, including leaf elemental content, spectral emissions, and root transcriptomics. By evaluating the combined and individual effects of soil type and root thermal treatments, we sought to reveal how external environmental cues propagate through microbial assemblages and ultimately reshape plant molecular and physiological states. This unified approach enables us to assess the response of the grapevine holobiont and more accurately capture its adaptive behaviour as a whole.

## 2. Materials and Methods

### 2.1 Experimental setup

One hundred cuttings of *Vitis vinifera* ‘Trebbiano Romagnolo’ clone R5 were grafted onto clonal rootstock Kober 5BB, rooted and grown under nursery conditions for one vegetative season at Vivai Cooperativi Rauscedo (Rauscedo, Italy), following standard practices of grapevine plant production. One-year-old cuttings were grown under open-field conditions at the experimental farm “Antonio Servadei” of the University of Udine from August 2021 (T0) until June 2022 (T1). Cuttings were planted in 2-litre potted soil using three commercially available growing substrates: (i) micronized river siliceous sand (Sabbia Fine Naturale, Axton; hereafter referred to as “sand”), (ii) mixture of sphagnum peat moss, green compost, and sand (Terriccio per Tappeti Erbosi, Geolia, Calvisano, Italy; hereafter referred to as “peat”), and (iii) sphagnum peat moss combined with bovine and equine manure (Terriccio con Stallatico, Geolia, Calvisano, Italy; hereafter referred to as “manure”). Half of the soil was subjected to a soil thermal treatment (i.e., autoclaved at 121 °C for 60 minutes) the day before planting. Half of the cuttings underwent root heat treatment, as follows. Heat treatment was applied by submerging roots in hot water at 50 °C for 45 minutes. The overall experimental design included 12 treatments: three soil types (3 levels: sand, peat, manure), root thermal treatment (2 levels: presence or absence), and soil thermal treatment (2 levels: presence or absence). Plants were irrigated using a drip irrigation system and allowed to grow for 11 months. NPK fertilizers were supplied on 01/06/2021 and 10/06/2021.

### 2.2 Sample collection

Bulk soil samples (in triplicate) were collected at planting (time 0, T0) and stored at -20 °C for further analyses. At the end of the experiment (time 1, T1), rhizosphere soil, root tissue, and leaves were collected from three potted plants per condition, as described below. An exception to this sampling plan was represented by the condition “sand autoclaved, root not treated”, for which only one plant was available, and for the conditions “manure non-autoclaved, root not treated” and “peat non-autoclaved, root not treated,” four samples were collected per condition. The complete list of samples is provided as **Supplementary File S1**. At T1, plants were uprooted, and soil was separated from the root ball using the pull and shake method to collect the rhizosphere and cleanse the roots for sampling (Duineveld *et al*., 1998). Rhizosphere samples were stored in 15 mL Falcon^®^ tubes on dry ice until their transfer at -80 °C. The remaining soil was collected and stored at 4 °C for chemical analyses. Apical portions of non-lignified roots were also collected, snap frozen in liquid nitrogen and stored in dry ice until storage at -80 °C. For elemental analysis, the leaves were sliced using a ceramic knife, placed in 50 mL Falcon tubes, and stored in dry ice until storage at -20°C.

### 2.3 Soil Analysis

Bulk soil obtained at T0 and non-root-associated soil obtained at T1 were homogenized using a 2 mm sieve. Soil samples were divided into aliquots: about 500 g of soil was weighed into an aluminium tray and air-dried for subsequent physico-chemical analyses, and 200 g of fresh soil were stored in plastic bags at 4 °C for subsequent ATP (Adenosine Triphosphate) quantification. Soil pH was measured in soil-water suspension (1:2.5, weight-to-volume ratio) using a pH-meter (Crison, Italy). Electrical conductivity (EC) was measured in a 1:2 soil-to-water ratio (Rhoades, 1996) using an XS Cond 7 Vio conductivity meter (Giorgio Bormarc s.r.l., Italy) and reported as µS/cm. The analysis of Dissolved Organic Matter (DOM) included the determination of dissolved organic carbon (DOC) and dissolved total nitrogen (DN), measured as milligrams of dissolved element per gram of soil. An aliquot of 4 g of dry soil was mixed with 40 mL of Milli-Q^®^ water and shaken overnight in the dark. After centrifugation and filtration, DOC and DN were measured using a TOC analyser (TOC-V_CPN_ Shimadzu, Kyoto). Total organic C and total N were measured on solid soil samples, after ball-mill grinding, with a CHNS elemental analyser (Vario Microcube, Elementar^©^). For organic C measurement, samples were previously treated with HCl to remove the inorganic C fraction. The C/N ratio was then calculated. The ATP extraction from fresh soil samples followed the acid-extraction method with modifications to account for ATP adsorption by soil colloids. Briefly, the extraction was carried out with trichloroacetic acid, orthophosphate, and imidazole solution (Redmile-Gordon, White and Brookes, 2011) with 2 min of sonification on an ice bath. An ATP spike was added as an internal standard to estimate its extraction recovery. The measurement of ATP in soil extracts was carried out using an ATP kit based on bioluminescence reaction (BioThema luminescent assay, Sweden) at a “Spark” multimode microplate reader (Tecan Trading AG, Switzerland). ATP is reported as nanomoles of ATP per gram of soil. Collinearity between pairs of traits was defined as a Pearson R^2^ > 0.9 at T0 and T1. In case of a collinear pair of traits, only one of the two was retained for analysis. The final physico-chemical parameters were DOC (Dissolved Organic Carbon), pH, ATP, and C/N.

### 2.4 Multispectral imaging of plants

At the end of the experiment, multispectral pictures of each plant were taken using two Micasense RedEdge-MX DUAL cameras. Images were captured by placing the potted plants on a flat surface against a neutral background, maintaining a consistent distance from the cameras. Before each image acquisition, a picture of the calibrated reflectance panel (CRP) was taken. CRP is a Lambertian surface with a reflectance calibration curve associated that allows for the conversion of raw pixel values into absolute reflectance. Moreover, to minimize the error during image capture due to changes in the light, a downwelling light sensor (DLS) has been coupled to the multispectral camera to adjust the readings to ambient light automatically. Since images captured at different wavelengths can have slight misalignments due to sensor differences and lens distortions, an image alignment step was performed before extracting reflectance values. This was done using the OpenCV package in Python, applying an affine transformation to correct for shifts, rotations, and scale variations between images. The transformation was computed using key point detection and feature matching across the different wavelength bands, ensuring accurate pixel-to-pixel correspondence. The wavelength values for each plant were extracted using ImageJ software by selecting an area on a leaf surface perpendicular to the image captured, leaf by leaf, with at least 100 pixels and at least two leaves per plant, and using the R package “raster”. This study focuses on NDVI (Normalized Difference Vegetation Index), CVI (Chlorophyll Vegetation Index), GDVI (Green Difference Vegetation Index), and NDWI (Normalized Difference Water Index). These indices use different waveband intensities to provide information on chlorophyll and water content, thus acting as proxies of plant physiology. Collinearity between pairs of traits was defined as a Pearson R^2^ > 0.9. No collinearity was observed.

### 2.5 Multielemental analysis of leaves

The concentration (parts per million, ppm) of macro-nutrients, Ca (calcium), K (potassium), Mg (magnesium), P (phosphorus), S (sulphur), and micro-nutrients, Cu (copper), Fe (iron), Mn (manganese), Na (sodium), Zn (zinc) in grapevine leaves was determined by Inductively Coupled Plasma–Optical Emission Spectroscopy (ICP-OES 5800, Agilent Technologies, Santa Clara, USA). Leaves were oven-dried for 72 h at 50-60 °C and ground into powder. For each sample, around 100 mg of ground powder was digested with concentrated ultrapure HNO3 using a microwave oven (ETHOS EASY, Milestone s.r.l., Sorisole, BG, Italy) according to the USEPA 3052 method “Plant Xpress” (Agency (USEPA), 1996). Element quantifications were carried out using certified multi-element standards. Collinearity between pairs of traits was defined as a Pearson R^2^ > 0.9. No collinearity was observed.

### 2.6 Nucleic acid extraction, library preparation, and sequencing

DNA was extracted from approximately 0.25 g of non-dried soil stored at -80 °C using the DNeasy PowerLyzer PowerSoil kit from QIAGEN (Hilden, Germany), following the manufacturer’s instructions. DNA was then sent to IGA Technology Services s.r.l. (Udine, Italy) for library preparation and sequencing. Libraries were prepared following the Illumina 16S Metagenomic Sequencing Library Preparation protocol in two amplification steps: an initial PCR amplification using locus-specific PCR primers and a subsequent amplification that integrates relevant flow-cell binding domains and unique indices (NexteraXT Index Kit, FC-131-1001/FC-131-1002). The locus-specific primers for the 16S rRNA gene were 341F (5’-CCTACGGGNGGCWGCAG-3’) and 805R (5’-GACTACHVGGGTATCTAATCC -3’). The locus-specific primers for the ITS rRNA were ITS3F (5’-GCATCGATGAAGAACGCAGC-3’) and ITS4R (5’-TCCTCCGCTTATTGATATGC-3’). Libraries were sequenced on NovaSeq 6000 instrument (Illumina, San Diego, CA) using a 250-bp paired-end mode.

For RNA sequencing, 100 mg of root samples were cleansed with tap water and ground in liquid nitrogen. Total RNA was extracted using the Spectrum Plant Total RNA Kit (SIGMA, The Netherlands). Libraries were prepared using the Universal Plus mRNA-Seq kit (Tecan Genomics, Redwood City, CA) following the manufacturer’s instructions. Libraries were sequenced on a NovaSeq6000 instrument (Illumina, San Diego, CA) using a 150-bp paired-end sequencing mode. Raw transcriptomic reads were deposited in GEO under the accession GSE287854. Raw metabarcoding reads were deposited in the NCBI Sequence Read Archive under the accession PRJNA1217140.

### 2.7 Data analysis

Adapters were removed using the *filterandtrim* function of DADA2 (Callahan *et al*., 2016). Classification of 16S rRNA and ITS sequences was performed using DADA2 against the SILVA (Quast *et al*., 2013) and UNITE (Kõljalg *et al*., 2020) databases, respectively. Rarefaction curves were plotted using the *vegan* R package (Oksanen *et al*., 2024). Amplicon sequence variants (ASVs) not classified at the Kingdom level as “Bacteria” or “Fungi” were removed. Raw ASVs’ counts were normalized by calculating their percentage relative to the total reads sequenced in each sample. ASVs with an average abundance lower than 0.001% were removed. Additional analyses were performed at the phylum and genus levels, aggregating ASVs based on their taxonomic classifications. The R (R Core Team, 2021) package *DESeq2* (Love *et al*., 2014) was utilized to perform differential abundance analysis. For this step, the ASVs were merged, considering the lowest taxonomical level reached by the single ASV. Non-Metric Multidimensional Scaling (NMDS) analysis and alpha diversity analysis were conducted using the *vegan* package (Oksanen *et al*., 2024). The variable factors were represented as vectors on the NMDS plot using the *envfit* function of the package *vegan*. Nested PERMANOVA analysis was performed using the *adonis* function of the package *vegan*. Functional prediction of bacterial species was performed using the FAPROTAX database and software (Louca *et al*., 2016). Fungal genera were metabolically classified using FUNGuild (Version 1.0) and a manually curated version of the FUNGuild database (Nguyen *et al*., 2016). The list of FAPROTAX and FUNGuild functional categories considered in this study is visible in **Supplementary Table S1**.

RNA-seq reads were aligned using STAR software (Dobin *et al*., 2013) against the *Vitis vinifera* 12Xv0 genome assembly of the strain PN40024 (GCA_000003745.2) using the V2.1 version of gene model prediction. *DESeq2* (Love, Huber and Anders, 2014) was employed for differential gene expression analysis. To associate functions to gene models, the expression table was compared with the Phytozome database (Goodstein *et al*., 2012) file “Vvinifera_457_v2.1.annotation_info.txt,” which includes the “Arabi_defline” column. This column provides the *Arabidopsis thaliana* gene description or annotation corresponding to the gene under analysis. GO annotation was performed by retrieving GO terms and their descriptors using the GO.db R package (Carlson, 2023) and the enrichR package (Kuleshov *et al*., 2016) for functional enrichment analysis. Classification of RNA viruses based on alignment of RNA sequences was performed using a previously described pipeline (Di Gaspero *et al*., 2022).

Data were plotted using the *pheatmap* (Kolde, 2019), *ggplot2* (Wilkinson, 2011), and *corrplot* (Wei *et al*., 2017) packages. Where not specified, the *cor.test* function from the *stats* package was used to perform Spearman correlation tests; P-values were adjusted for multiple testing using the false discovery rate (FDR) correction method. All scripts and R functions used in this study are available at https://github.com/genomeud/intevine.

## 3. Results

### 3.1 Physico-chemical parameters of soil

The physico-chemical parameters retained after collinearity analysis were DOC, pH, ATP and C/N (Supplementary Tables S2, S3). The four parameters showed significant differences across soil types, both at T0 and T1 (Supplementary Figs. S1 and S2, respectively). Such variations defined three different clusters in the ordination space of the NMDS plot at the two sampling points (Supplementary Fig. S3, Fig. 1a;), with soil type explaining up to 89% and 86% of the variability, respectively, according to PERMANOVA analysis (Supplementary Tables S4, S5). A moderate effect of the soil autoclave treatment was observed both at T0 and T1; treatment of the root before transplantation did not affect the physico-chemical properties of the soil at T1.

**Figure 1.**
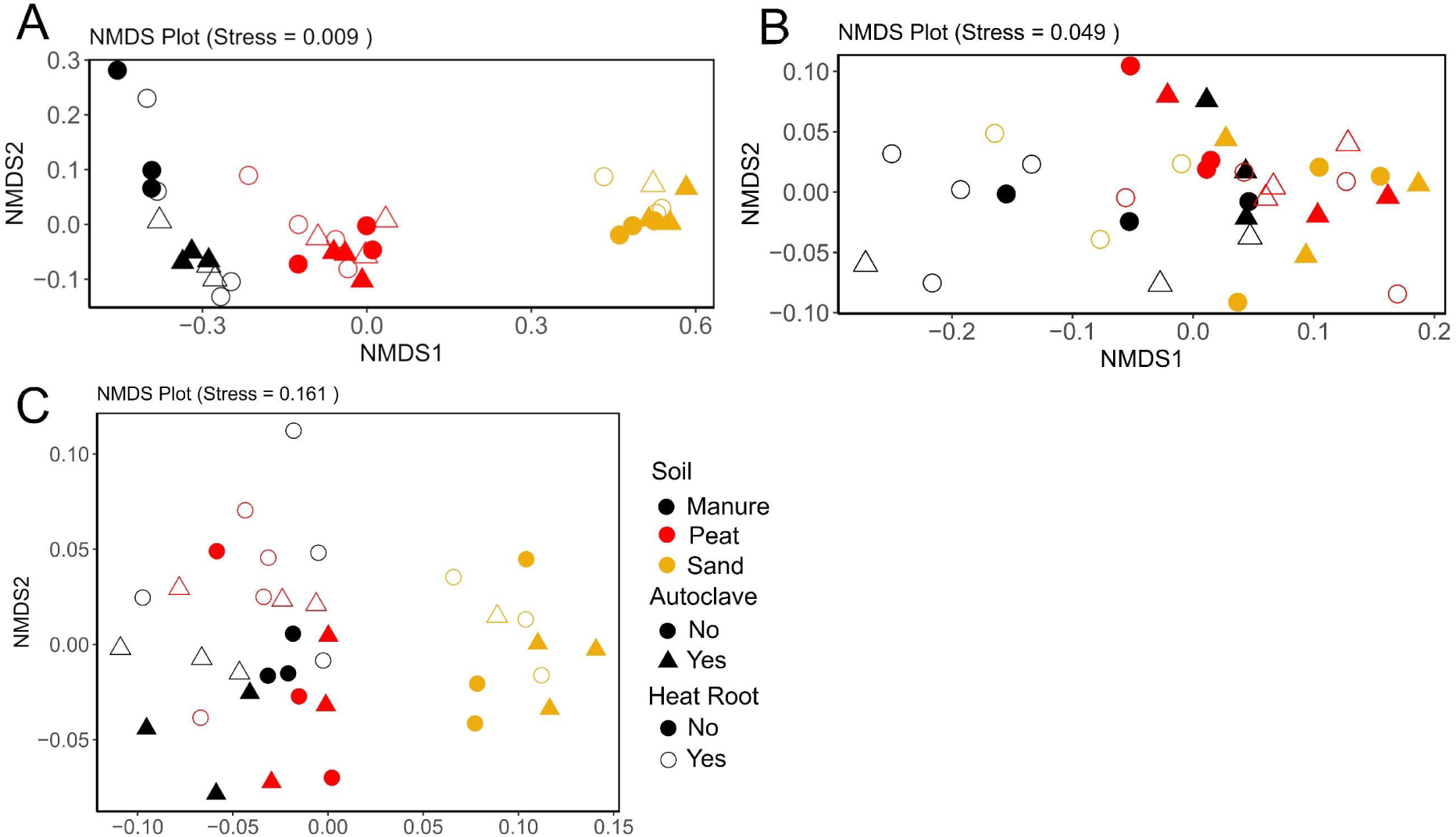
Ordination of soil chemistry and leaf traits in grapevines after growth in different soils. Non-metric multidimensional scaling (NMDS) plot illustrating separation among (**A**) soil samples by chemical composition, (**B**) leaf samples by multispectral imaging indices, and (**C**) leaf samples by multielemental profiles at the end of the experiment (T1, 18 months).

### 3.2 Multispectral indices

Multispectral indices from collected leaves did not show a specific pattern of distribution in the NMDS ordination (**Fig. 1B**) as observed for soil physico-chemical analysis. However, soil type and the heat treatment of roots within soil-autoclave treatments significantly influenced the multispectral indices, explaining 27% and 23% of the variance, respectively, according to nested PERMANOVA analysis (**Supplementary Table S6**). Among the indices evaluated, only CVI and NDWI showed significant differences between soil types or heat root treatment (**Supplementary Figs. S4** and **S5**), while none of the indices were affected by soil autoclaving.

### 3.3 Multi-element analysis

NMDS analysis based on ion concentrations in the leaves separated sand samples from peat and manure (Fig. 1c). Nested PERMANOVA analysis showed that soil type was the only factor significantly contributing to this differentiation pattern (**Supplementary Table S7**). Such diversity is driven by seven of the analysed elements that differed significantly in concentration across the three soil types (**Supplementary Fig. S6**)**3.4 Soil bacterial microbiome.** A total of 3,794,673 and 24,597,687 bacterial reads were retained for samples collected at T0 and T1, respectively (summary statistics in **Supplementary Table S8**), providing sufficient sequencing depth to ensure robust coverage of the bacterial community diversity (**Supplementary Fig. S7A,B**). At T0, diversity was shown only for the peat and manure samples, as the DNA recovered from the sand samples was not successfully amplified, likely due to the low abundance of microorganisms in this substrate before plant growth and irrigation. At the beginning of the experiment, peat soil showed a higher relative abundance of *Spirochaetota* and *Actinobacteriota* compared to manure soil, with *Gemmatimonadota* also being more abundant in peat soil (**Supplementary Fig. S8A**). *Bacillota* exhibited a higher relative abundance in autoclaved soil than in non-autoclaved soil, while *Proteobacteriota, Verrucomicrobiota*, and *Bacteroidota* were more abundant in non-autoclaved soil. NMDS analysis performed on bacterial ASVs showed that samples clustered by soil type and soil thermal treatment, defining significantly different groups according to PERMANOVA analysis (Supplementary Fig. S8B, Table S9). No significant differences in species richness and Shannon’s index were observed (Supplementary Fig. S8D,E).

The differences in microbiome composition across soil types persisted in the rhizosphere and adjacent soil (i.e., root surrounding soil) at T1 (**Fig. 2**; PERMANOVA, **Supplementary Table S10**). While the most abundant phyla did not vary across soil (**Fig. 2A**), cluster analysis and NMDS revealed a separation of soils into three main clusters based on soil type; each cluster is further divided into two sub-clusters based on soil thermal treatment (i.e., autoclaving) (**Fig. 2B,C**, **Supplementary Fig. S9**). In addition, even if to a lesser extent, root thermal treatment influences the assembly of bacterial communities (**Supplementary Fig. S**9).Soil physico-chemical parameters shown association with the bacterial communities. **Fig. 2C** shows NMDS of samples based on soil bacterial microbiome together with vectors of physico-chemical variables significantly affecting the NMDS ordination. C/N ratio was higher in peat, pH was higher in sand, and DOC and ATP were higher in manure.

**Figure 2.**
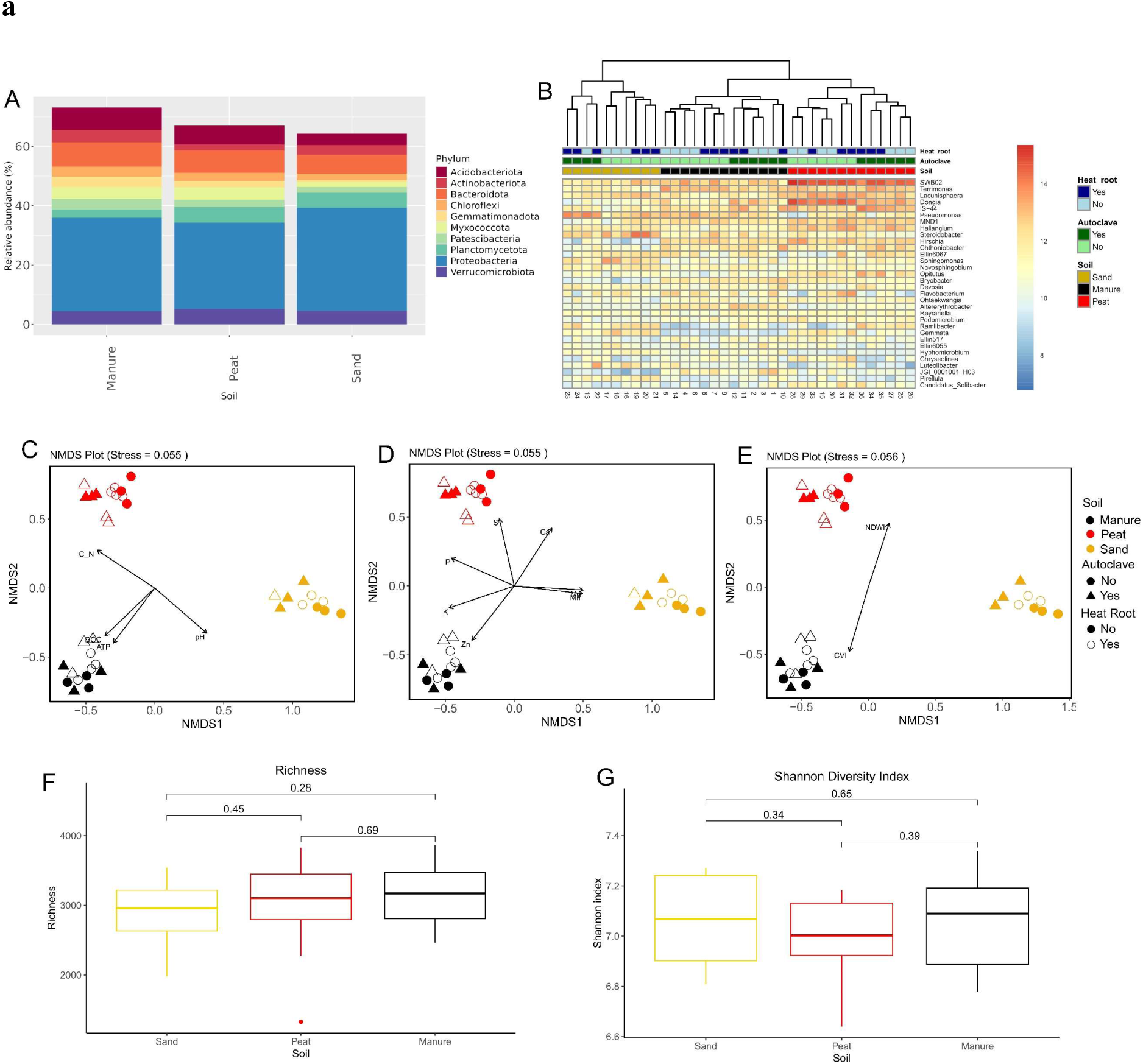
Rhizosphere bacterial microbiome associated with grapevine after growth in different soils. (**A**) Average relative abundance of the most predominant phyla. (**B**) Heatmap of the most abundant genera with clustering analysis. NMDS analysis of the T1 bacterial rhizosphere microbiome, combined with (**C**) physico-chemical variables, (**D**) leaf multielemental concentrations, and (**E**) multispectral imaging indices, includes vectors representing only significant correlations identified by the envfit function. Boxplot with pairwise-Wilcoxon test (in brackets) of the (**F**) richness (i.e., observed number of ASVs; Kruskal-Wallis, P = 0.5) and (**G**) Shannon index (Kruskal-Wallis, P = 0.51) across the three types of soil.

Relationship were also found between the bacterial community and the multielemental analysis (**Fig. 2D)** between NMDS coordinates and nutrient concentration in leaves. Manure soil shows higher levels of potassium and zinc, peat shows higher level of sulphur and phosphorus, and sand shows higher levels of magnesium

**Fig. 2E** shows the relationship between NMDS coordinates and multispectral imaging. Since from one of the sand samples (sample 13) it was not possible to extract the wavelength data, the NMDS was performed without it, hence the differences if compared with the previous. The manure soil cluster is associated with higher CVI and peat is associated with higher NDWI. The three soil types did not show significant differences in alpha diversity measured as species richness and Shannon’s diversity index (**Fig. 2F,G**). However, the composition of individual taxa showed several differences. A total of 542 bacterial genera exhibited differential abundance in at least one comparison, including soil type contrasts, soil sterilization treatments, or root heat treatments. (**Supplementary File S2**). Seventeen bacterial genera showed different abundances across all soil types, both at T0 and T1. Among them are *Spirochaeta*, *Anaerolinea*, *Dongia*, and *Haliangium*. The effect of autoclaving soil resulted in 50 differential genera at T0 and 26 at T1. Only two of them, *Bacillus* and *Verrucosispora*, showed differences in both time points. Root thermal treatment had a minor effect on bacterial communities. Only three genera showed differences as a consequence of the treatment: *Caedibacter*, *Desulfuromonas*, and the I-8 genus of the family *Phycisphaeraceae*.

### 3.5 Soil fungal microbiome

A total of 2,605,034 and 8,482,398 fungal reads were generated from ITS amplicons at T0 and T1, respectively. These reads were assigned to 600 and 774 fungal ASVs, respectively (Summary statistics are presented in **Supplementary Table S8**). Before planting (T0), peat samples were dominated by *Ascomycota,* and *Basidiomycota* abundance was negligible, while manure showed some contribution of *Basidiomycota* to the total fungal community (Supplementary Fig. S10A). NMDS, clustering and PERMANOVA analysis reveal the presence of two distinct clusters corresponding to the soil types, with the autoclave treatment further differentiating within each soil type (**Supplementary Fig. S10B,C; Supplementary Table S11**). Different soils showed different levels of fungal diversity; peat showed higher values of richness and Shannon’s index than manure (**Supplementary Fig. S10D,E**). At T1, the soil retained a high relative abundance of *Ascomycota* and *Basidiomycota*. In addition, *Glomeromycota* were present at relatively high abundance, compared to what observed at T0 (**Fig 3A**). Clustering analysis based on abundance of fungal genera showed some clustering based on soil type and some clusters determined by root heat treatment (**Fig. 3B**). NMDS analysis showed limited clustering across soil type and limited or no clustering across conditions (**Fig 3C-E**), although PERMANOVA analysis indicated substantial contribution of all conditions to the variance (Supplementary Table S12). Higher DOC and ATP levels were associated with fungal communities in manure soil clusters, pH with those in sand, and C/N with peat ones (**Fig 3C**). Increased levels of manganese, calcium, and magnesium were observed in the sand cluster, and a higher concentration of potassium in the manure cluster (**Fig. 3D**). Fungal communities showed significant correlations with the same multispectral indices as the bacterial population: the manure cluster was associated with higher CVI values and peat with higher NDWI values (**Fig. 3E**). When performing NMDS separately in each soil type, no major effect of soil thermal treatment or soil autoclaving was observed in each soil (**Supplementary Fig. S11**). However, several individual taxa showed different abundance across conditions and treatments. Specifically, a total of 84 fungal genera showed differences in at least one comparison between soil types (**Supplementary File S3**). Only the genus *Kernia* showed differences across all soil types, and only the genus *Chetomium* showed different abundance as a consequence of autoclaving soil at both time points. Ten genera showing differential abundance after thermal root treatment, namely *Qarounispora, Rhizoglomus, Septoglomus, Oliveonia, Glomus, Fusarium, Clonostachys, Amniculicola, Lophiostoma* and *Trichomonascus*. Richness and Shannon index were higher in peat, intermediate in sand and lower in manure soil (**Fig. 3F,G**).

**Figure 3.**
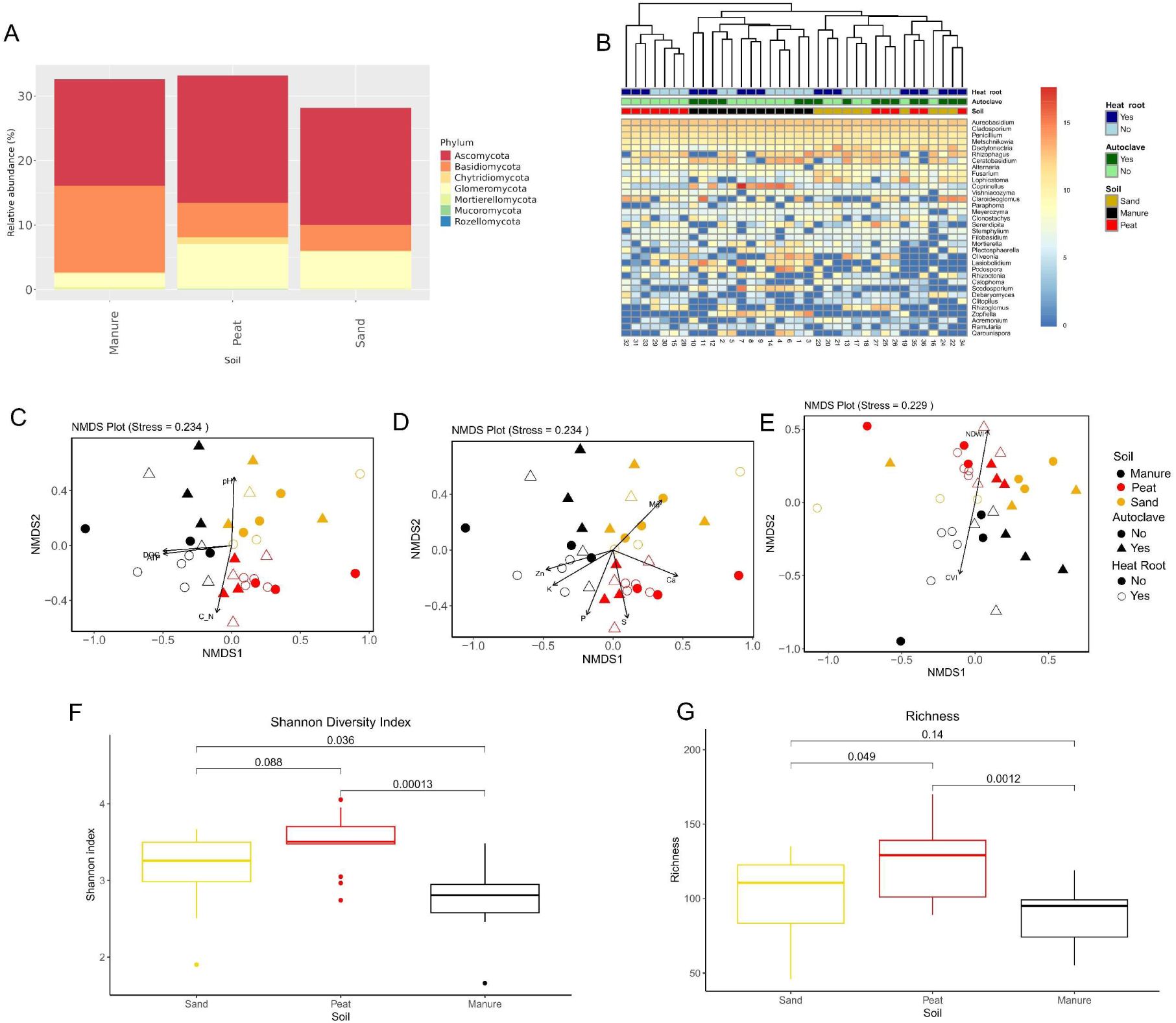
Rhizosphere fungal microbiome associated with grapevine after growth in different soils. (**A**) Average relative abundance of the phyla identified. (**B**) Heatmap of the most abundant genera with clustering analysis. NMDS analysis of the T1 fungal rhizosphere microbiome, combined with (**C**) physico-chemical variables, (**D**) leaf multielemental concentrations, and (**E**) multispectral imaging indices, includes vectors representing only significant correlations identified by the envfit function, highlighting relationships between these parameters and bacterial distribution. Boxplot with pairwise-Wilcoxon test (in brackets) of the (**F**) richness (i.e., number of observed fungal ASVs; Kruskal-Wallis, P = 0.00089) and (**G**) Shannon index (Kruskal-Wallis, P = 0.0029) in the three types of soil.

### 3.6 Functional prediction of bacterial and fungal taxa

Functional prediction based on the taxonomic affiliation of bacterial and fungal clades allowed the identification of categories related to plant-association as mutualistic or pathogen functional traits (**Supplementary Table S1**). The distribution of such categories revealed that potential plant pathogens were less abundant in peat compared to other soils (**Fig. 4A**). Bacteria involved in processes related to the removal of nitrogen from the soil reservoir, such as nitrogen and nitrate respiration showed the higher relative abundance in sand and the lower in peat (**Fig. 4B**), while those able to perform nitrogen fixation were more abundant in sand and nearly absent in manure and peat (**Fig. 4C**). The potential fungal phytopathogens was significantly less abundant in the root surrounding soil of plants grown in manure soil compared to those from the other two types of soils (**Fig. 5A**). Fungi classified by FUNGUILD as arbuscular mycorrhizae did not show significant variation across soil types (**Fig. 5B**). Fungal parasites were more abundant in peat than in sand and manure (**Fig. 5C**). Finally, endophytic fungi (**Fig. 5D**) showed lower values in sand compared to peat, and no significant differences in other comparisons.

**Figure 4.**
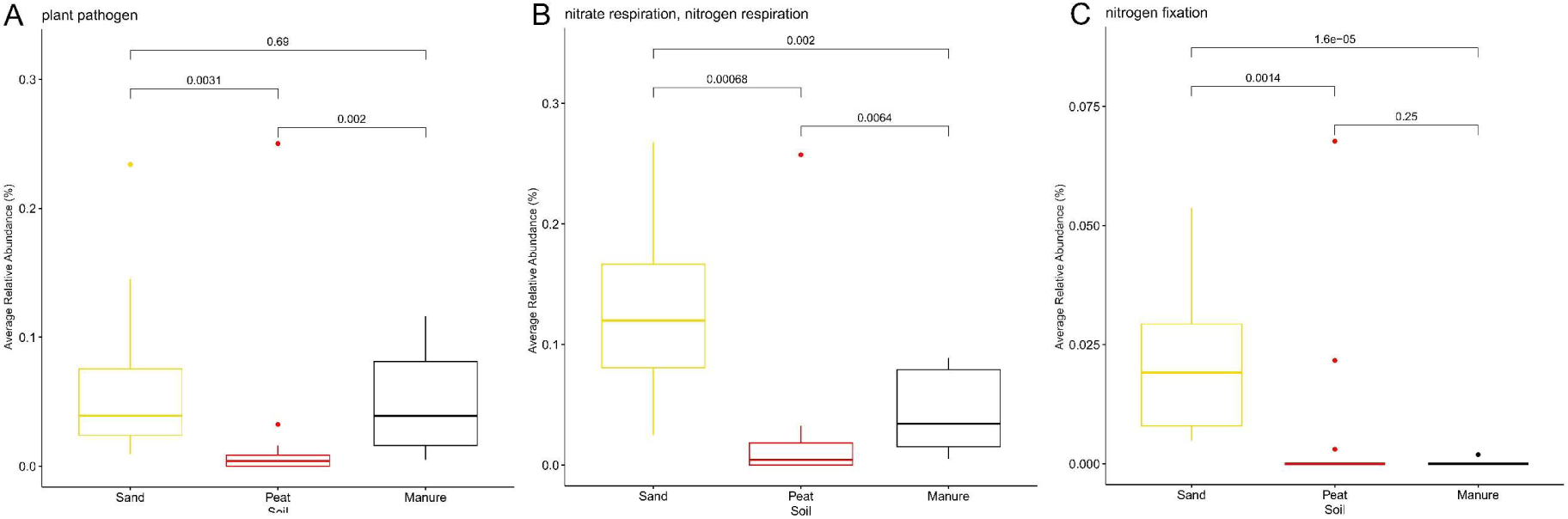
Effect of soil in shaping predicted bacterial functional groups. Boxplot with pairwise-Wilcoxon test (in brackets) comparing the relative abundance of functional classes of bacterial species across the three types of soil at T1. Bacterial species related to (**A**) plant pathogens (Kruskal−Wallis, P = 0.0016), (**B**) nitrogen and nitrate respiration (Kruskal−Wallis, P = 0.00014), and (**C**) nitrogen fixation (Kruskal−Wallis, P = 0.000015).

**Figure 5.**
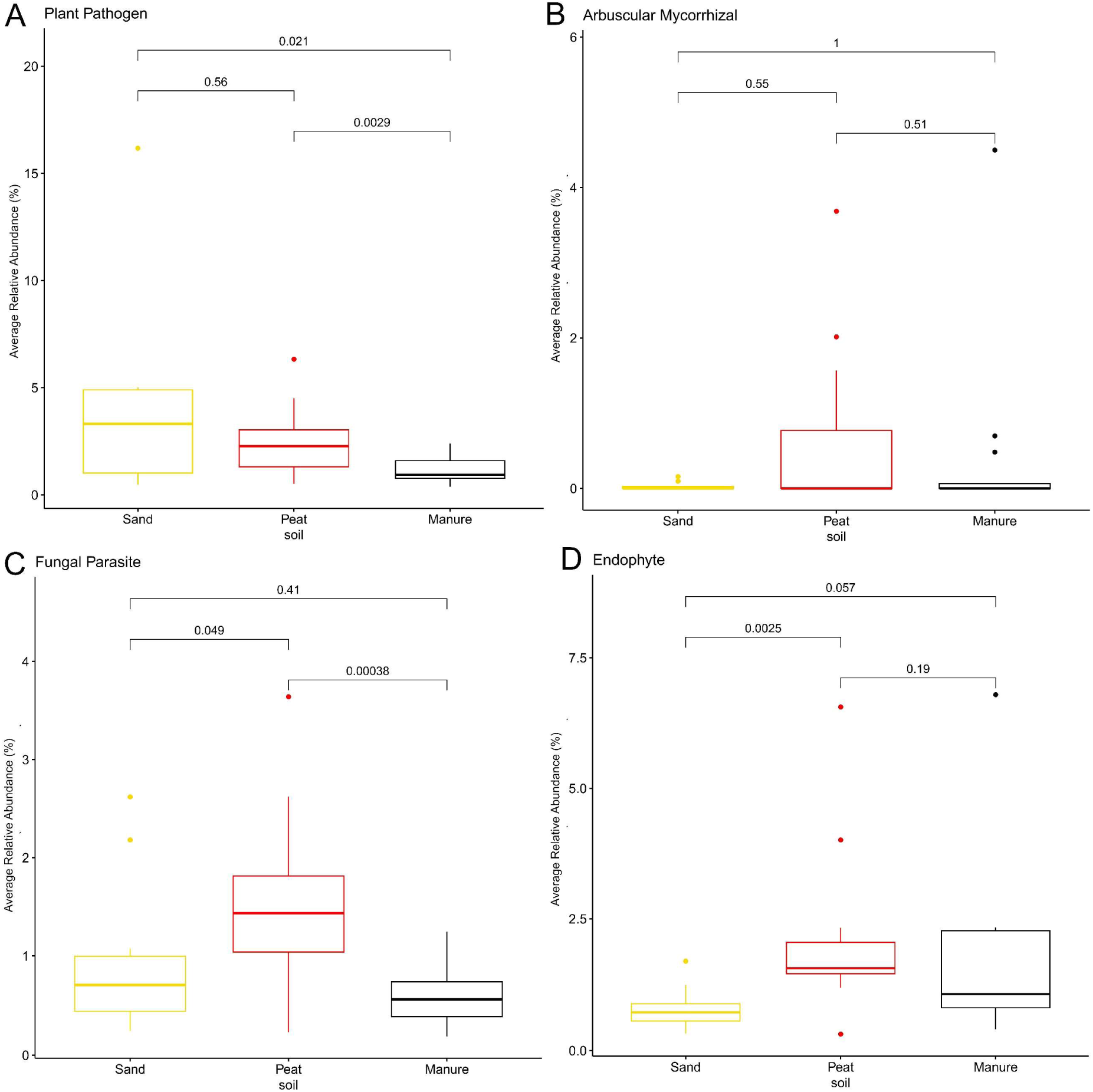
Effect of soil in shaping predicted fungal functional guilds. Boxplot with pairwise-Wilcoxon test comparing the relative abundance of functional classes of fungal genera across the three types of soil at T1. Fungal genera classified as (**A**) plant pathogens (Kruskal−Wallis, P = 0.0085), (**B**) arbuscular mycorrhizal (Kruskal−Wallis, P = 0.0031), (**C**) fungal parasites (Kruskal−Wallis, P = 0.0014), and (**D**) endophyte mycorrhizal (Kruskal−Wallis, P = 5e-04).

### 3.7 Transcriptome analysis

On average, 21.8 million fragments per sample were properly mapped, ranging from 5.4 million to 48 million. Clustering analysis (**Supplementary Fig. S12**) and NMDS (**Supplementary Fig. S13**) did not show clear separation between soil types and conditions. PERMANOVA analysis shows that soil types, soil sterilization, and root thermal treatment are significantly affecting the root transcriptome, although with a relatively small proportion of explained variance (**Supplementary Table S13**). Differential expression analysis identified 3,489 differentially expressed genes (DEGs) when comparing expression in peat with expression in manure (peat vs. manure), 65 in sand vs. peat, and 5,467 in sand vs. manure. Additionally, 1,116 DEGs were identified in response to root heat treatment, while no DEGs were observed in response to autoclave treatment of soil substrates (**Supplementary File S4**). Genes differentially expressed in at least one soil comparison were enriched for GO categories related to photosynthetic activity, including chloroplast stroma and lipid metabolic activity (**Fig. 6A**). When assessing the effect of root thermal treatment, DEGs were enriched in terms related to translation processes and ribosome formation (**Fig. 6B**).

**Figure 6.**
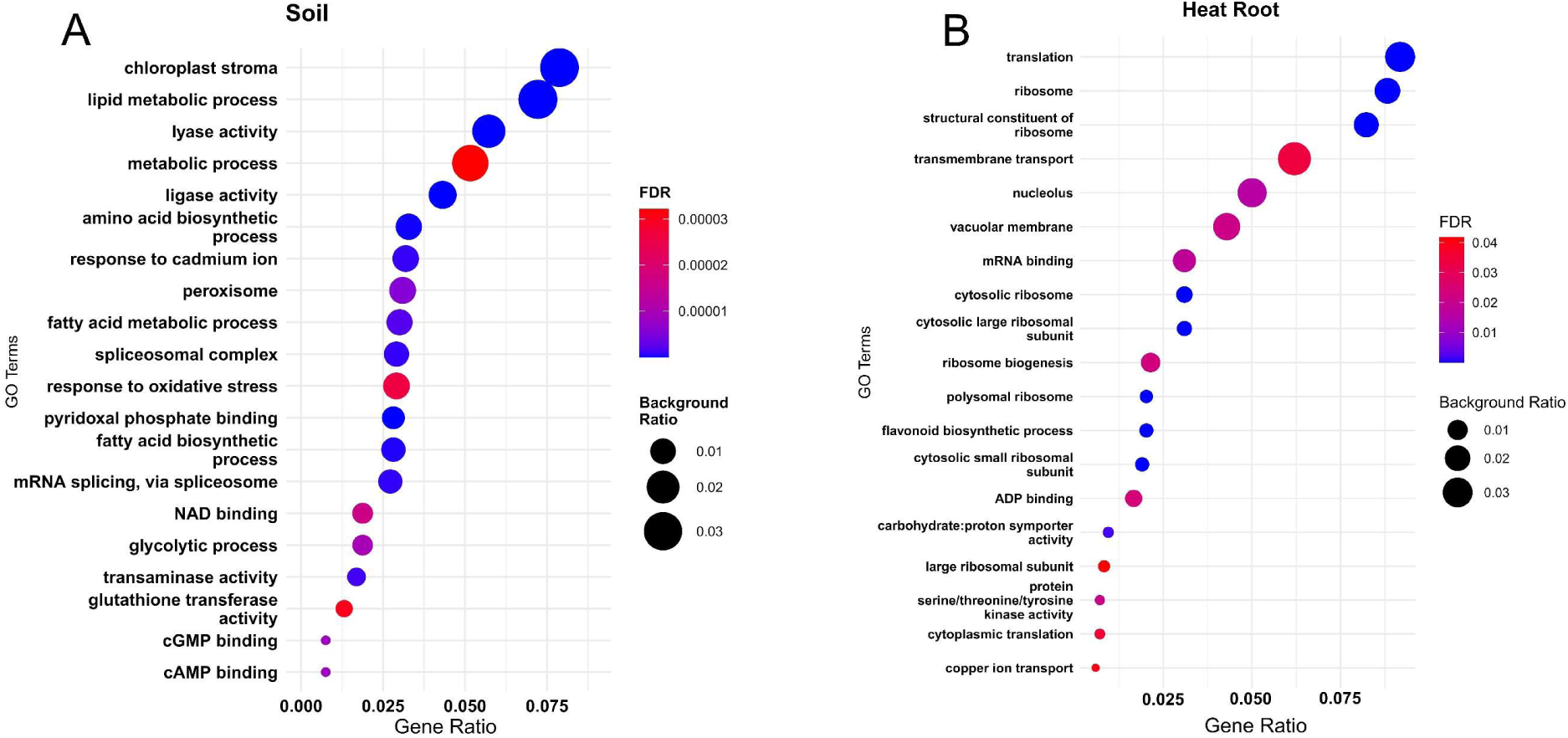
Differential gene expression in grapevine root tissues under different conditions. Gene ontology (GO) enrichment dot plot illustrating functional enrichment analysis. Gene ratio represents the proportion of genes associated with each GO term, background ratio indicates the proportion of genes in the reference set, and False Discovery Rate (FDR) reflects statistical significance. (A) Top 20 GO categories differentially enriched in at least one soil comparison. (B) All differentially enriched GO categories in the heat root treatment

RNA-seq also enabled the detection of viral RNA; our results revealed the presence of four viruses, namely *Grapevine Pinot Gris Virus, Grapevine Rupestris Stem-pitting associated virus, Grapevine yellow-speckle viroid and hop stunt viroid*. Viral abundance was partly influenced by soil type, as evidenced by clustering analysis (**Supplementary Fig. S14A**). **Supplementary Fig. S14B** shows total viral abundance as a function of DOC, stratified by soil type and soil conditions. A positive correlation was observed between DOC and total viral abundance (r = 0.3692, P = 8.491e-05). In addition, Wilcoxon test showed that viral abundance was higher in manure **(Supplementary Fig. S14C**).

## 4. Discussion

We present an integrative analysis of multi-omics framework to dissect how soil type and root thermal treatments shape the grapevine holobiont. By combining soil physico-chemical features, leaf ionomic measurements, multispectral imaging, root surrounding soil bacterial and fungal surveys, and root transcriptomics, we captured cross-scale linkages from edaphic inputs through microbial assembly to host physiology. In doing so, we showed that soil and its chemistry act as the master regulators: distinct soil types not only harboured unique bacterial and fungal communities but also imposed their signature on leaf nutrient profiles and spectral patterns, as well as shifts in root gene expression. Such a holistic perspective not only illuminates the mechanistic foundations of grapevine adaptation to variable environments but also provides actionable insights for optimizing vineyard management.

Grapevine is cultivated worldwide across a wide range of soil types, from sandy to clay-rich and loamy soils (Deloire *et al*., 2005; Uyan, Janus and Ertunç, 2023). Differences in soil properties influence water retention, nutrient availability, and root development, all of which affect grapevine performance and physiology (Quezada *et al*., 2014; Wang *et al.,* 2015). Moreover, in line with the plant-soil-microbe feedback (Bettenfeld *et al*., 2022), soil properties also shape the composition and functional potential of edaphic microbiomes, including those associated with the rhizosphere and plants (Darriaut *et al*., 2022). A key factor is the type and amount of organic carbon present in the soil, which affects microbial activity and nutrient cycling (Hu *et al*., 2021). Soils rich in stable organic matter, such as humified carbon, tend to promote microbial diversity and activity (Wang *et al*., 2025), as well as healthier vines (Visconti *et al*., 2024). Our data reinforced this paradigm and showed how bacterial community composition emerged as the most sensitive indicator of soil-driven grapevine holobiont differentiation under when grown under different soils, even for a relative short time (11 months). Although soil type wielded the strongest influence, autoclaving treatment reveals that artificial “sterilization” can nonetheless leave a lasting imprint on the microbiome. Soil autoclaving is experimentally used for biotic clearing of soils, alters microbial composition, and is used to foster microbial recolonization (King *et al*., 2024). While the effect of this treatment disrupted microbial communities at the initial stage of the experiment, a swift recovery of community assemblages was observed 11 months post-treatment, suggesting high resilience or rapid recolonization. However, such effect showed different levels between bacterial and fungal communities, where the former maintains a distinct trajectory, possibly due to higher sensitivity to initial treatment and/or a more defined niche colonization/selection and less heterogeneity across samples that allow the detection of a significant differentiation pattern. This persistent bacterial shift mirrors shorter-term legacies of soil treatments observed in vegetable and cereal systems (Li et al. 2019; DiLegge, Manter & Vivanco 2022), underscoring autoclaving as an enduring perturbation that reshapes bacterial communities, especially when external microbial inputs are limited.

Even if to a different extent, thermal root treatment mediates a shift in the fungal community assembly. Root thermal treatment is indeed a mainstay in grapevine nurseries to eliminate phylloxera, phytoplasmas, and other vascular pathogens from cuttings (Waite and May, 2005; Gramaje and Armengol, 2011; Clarke *et al*., 2019; Laimer and Bertaccini, 2019), however the collateral impacts on the native root-soil continuum associated microbiota that will be further established when grapevine cuttings are transplanted have rarely been examined. By stripping away resident symbionts and pathogens alike, root-heating treatments likely act as a strong ecological “reset” at the root–soil interface. Thermal shocks create a temporal vacuum in which the first post-treatment colonizers can access exudates and attachment sites (Eichmeier *et al*., 2018; Li *et al*., 2022), thus reshaping successional trajectories, with rapidly recolonizing species gaining priority. As expected, given that the new colonizers are drawn entirely from the bulk soil pool (here as sand, peat, and manure, autoclaved or not) (Zarraonaindia *et al*., 2015), it is plausible to expect that such treatment has a lower impact on the microbial community when different soils are compared. In fact, differences were observed only when the soil was analysed separately. Over time, such shifts in microbial colonizers of rhizosphere and root surrounding soil can influence nutrient cycling, hormone signalling, and defence priming, potentially affecting root architecture and physiology, as well as aboveground growth (Wankhade *et al*., 2025). For instance, our dataset revealed that over a thousand root transcripts remained differentially expressed a season later.

The findings of this study highlight how soil type influences the grapevine-microbiome holobiont as a whole, affecting plant physiology, nutrient uptake, and microbial interactions. These results underscore the importance of considering the plasticity of soil microbiomes and the potential of targeted soil treatments to shape soil–microbe–plant interactions in favour of plant-soil-microbe positive feedback, i.e., enrichment of mutualistic interactions. Such knowledge can inform more sustainable and effective soil management strategies, both in grapevine nurseries and in vineyard management, ultimately supporting vine propagation and plant productivity.

## Acknowledgements

Authors gratefully acknowledge the help of Nicoletta Felice from the University of Udine for DNA and RNA extraction. Authors gratefully acknowledge the help of Romina Carpi from Azienda Sperimentale A. Servadei, for technical help in plant cultivation. We thank IGA technology services s.r.l. for support in sequencing experiments. We also thank Laura D’Andrea for her assistance with DNA extraction and soil chemistry analysis. The authors also wish to thank the VSRP Internship Program of King Abdullah University of Science and Technology (KAUST) for supporting this research.

## Author contributions

Massimo Guazzini: Software, Investigation, Data Curation, Writing - Original Draft, Writing - Review & Editing. Ramona Marasco: Investigation, Writing - Review & Editing, Supervision. Slobodanka Radovich: Investigation, Resources, Writing - Review & Editing. Elisa Pellegrini: Investigation, Resources, Writing - Review & Editing. Marco Vuerich: Investigation, Resources. Arianna Lodovici: Investigation, Writing - Review & Editing. Giorgia Dubsky De Wittenau: Investigation. Eleonora Paparelli: Investigation. Gabriele Magris: Investigation. Laura Zanin: Investigation, Resources. Marco Contin: Investigation, Resources, Writing - Review & Editing. Elisa De Luca: Resources, Writing – Review & Editing. Daniele Daffonchio: Resources, Writing - Review & Editing, Supervision. Gabriele Di Gaspero: Conceptualization, Investigation, Resources, Writing - Review & Editing. Fabio Marroni: Conceptualization, Software, Investigation, Resources, Supervision, Writing - Review & Editing.

## Data availability

Raw RNA-seq reads were deposited in GEO under the accession GSE287854. Raw metabarcoding reads were deposited in the NCBI Sequence Read Archive under the accession PRJNA1217140. Scripts and functions used for analysis are available at https://github.com/genomeud/intevine.

## Bibliography

Agency (USEPA), U.S.E.P. (1996) EPA Method 3052: Microwave Assisted Acid Digestion of Siliceous and Organically Based Matrices. U.S. Environmental Protection Agency. Available at: https://www.epa.gov.

Berendsen, R.L., Pieterse, C.M.J. and Bakker, P.A.H.M. (2012) ‘The rhizosphere microbiome and plant health’, Trends in Plant Science, 17(8), pp. 478–486. Available at: 10.1016/j.tplants.2012.04.001.

Berg, G. et al. (2024) ‘Shared governance in the plant holobiont and implications for one health’, FEMS Microbiology Ecology, 100(3), p. fiae004. Available at: 10.1093/femsec/fiae004.

Bettenfeld, P. et al. (2022) ‘The microbiota of the grapevine holobiont: A key component of plant health’, Journal of Advanced Research, 40, pp. 1–15. Available at: 10.1016/j.jare.2021.12.008.

Callahan, B.J. et al. (2016) ‘DADA2: High-resolution sample inference from Illumina amplicon data’, Nature Methods, 13(7), pp. 581–583. Available at: 10.1038/nmeth.3869.

Carlson, M. (2023) GO.db: A set of annotation maps describing the entire Gene Ontology.

Clarke, C. w., et al. (2019) ‘Hot water immersion as a disinfestation treatment for grapevine root cuttings against genetically diverse grape phylloxera Daktulosphaira vitifoliae Fitch’, Australian Journal of Grape and Wine Research, 25(4), pp. 396–403. Available at: 10.1111/ajgw.12407.

Darriaut, R. et al. (2022) ‘Soil composition and rootstock genotype drive the root associated microbial communities in young grapevines’, Frontiers in Microbiology, 13. Available at: 10.3389/fmicb.2022.1031064.

Deloire, A. et al. (2005) ‘Grapevine responses to terroir: A global approach’, Journal International des Sciences de la Vigne et du Vin, 39, pp. 149–162. Available at: 10.20870/oeno-one.2005.39.4.888.

Di Gaspero, G. et al. (2022) ‘Evaluation of sensitivity and specificity in RNA-Seq-based detection of grapevine viral pathogens’, Journal of Virological Methods, 300, p. 114383. Available at: 10.1016/j.jviromet.2021.114383.

Dobin, A. et al. (2013) ‘STAR: ultrafast universal RNA-seq aligner’, Bioinformatics (Oxford, England), 29(1), pp. 15–21. Available at: 10.1093/bioinformatics/bts635.

Duineveld, B.M. et al. (1998) ‘Analysis of the dynamics of bacterial communities in the rhizosphere of the chrysanthemum via denaturing gradient gel electrophoresis and substrate utilization patterns’, Applied and Environmental Microbiology, 64(12), pp. 4950–4957. Available at: 10.1128/AEM.64.12.4950-4957.1998.

Eichmeier, A. et al. (2018) ‘High-throughput amplicon sequencing-based analysis of active fungal communities inhabiting grapevine after hot-water treatments reveals unexpectedly high fungal diversity’, Fungal Ecology, 36, pp. 26–38. Available at: 10.1016/j.funeco.2018.07.011.

El-Ramady, H.R. et al. (2014) ‘Soil Quality and Plant Nutrition’, in H. Ozier-Lafontaine and M. Lesueur-Jannoyer (eds) Sustainable Agriculture Reviews 14: Agroecology and Global Change. Cham: Springer International Publishing, pp. 345–447. Available at: 10.1007/978-3-319-06016-3_11.

Ghatak, A. et al. (2022) ‘Root exudation of contrasting drought-stressed pearl millet genotypes conveys varying biological nitrification inhibition (BNI) activity’, Biology and Fertility of Soils, 58(3), pp. 291–306. Available at: 10.1007/s00374-021-01578-w.

Goodstein, D.M. et al. (2012) ‘Phytozome: a comparative platform for green plant genomics’, Nucleic Acids Research, 40(D1), pp. D1178–D1186. Available at: 10.1093/nar/gkr944.

Gramaje, D. et al. (2022) ‘Exploring the Temporal Dynamics of the Fungal Microbiome in Rootstocks, the Lesser-Known Half of the Grapevine Crop’, Journal of Fungi, 8(5), p. 421. Available at: 10.3390/jof8050421.

Gramaje, D. and Armengol, J. (2011) ‘Fungal Trunk Pathogens in the Grapevine Propagation Process: Potential Inoculum Sources, Detection, Identification, and Management Strategies’, Plant Disease, 95(9), pp. 1040–1055. Available at: 10.1094/PDIS-01-11-0025.

Hassani, M.A., Durán, P. and Hacquard, S. (2018) ‘Microbial interactions within the plant holobiont’, Microbiome, 6(1), p. 58. Available at: 10.1186/s40168-018-0445-0.

Herz, K. et al. (2018) ‘Linking root exudates to functional plant traits’, PloS One, 13(10), p. e0204128. Available at: 10.1371/journal.pone.0204128.

Hu, H. et al. (2021) ‘Significant association between soil dissolved organic matter and soil microbial communities following vegetation restoration in the Loess Plateau’, Ecological Engineering, 169, p. 106305. Available at: 10.1016/j.ecoleng.2021.106305.

Islam, W. et al. (2020) ‘Role of environmental factors in shaping the soil microbiome’, Environmental Science and Pollution Research, 27(33), pp. 41225–41247. Available at: 10.1007/s11356-020-10471-2.

King, W.L. et al. (2024) ‘Autoclaving is at least as efective as gamma irradiation for biotic clearing and intentional microbial recolonization of soil’, mSphere, 9(7), pp. e00476–24. Available at: 10.1128/msphere.00476-24.

Kolde, R. (2019) ‘Pheatmap: pretty heatmaps’, R package version, 1(2), p. 726.

Kõljalg, U. et al. (2020) ‘The Taxon Hypothesis Paradigm—On the Unambiguous Detection and Communication of Taxa’, Microorganisms, 8(12), p. 1910. Available at: 10.3390/microorganisms8121910.

Kuleshov, M.V. et al. (2016) ‘Enrichr: a comprehensive gene set enrichment analysis web server 2016 update’, Nucleic Acids Research, 44(Web Server issue), pp. W90–W97. Available at: 10.1093/nar/gkw377.

Lade, S.B. et al. (2022) ‘Hot Water Treatment Causes Lasting Alteration to the Grapevine (Vitis vinifera L.) Mycobiome and Reduces Pathogenic Species Causing Grapevine Trunk Diseases’, Journal of Fungi, 8(5), p. 485. Available at: 10.3390/jof8050485.

Lade, S.B., Štraus, D. and Oliva, J. (2022) ‘Variation in Fungal Community in Grapevine (Vitis vinifera) Nursery Stock Depends on Nursery, Variety and Rootstock’, *Journal of Fungi (Basel*, Switzerland), 8(1), p. 47. Available at: 10.3390/jof8010047.

Laimer, M. and Bertaccini, A. (2019) ‘Phytoplasma Elimination from Perennial Horticultural Crops’, in A. Bertaccini et al. (eds) Phytoplasmas: Plant Pathogenic Bacteria - II: Transmission and Management of Phytoplasma - Associated Diseases. Singapore: Springer, pp. 185–206. Available at: 10.1007/978-981-13-2832-9_9.

Lanyon, D., Cass, A. and Hansen, D. (2004) ‘The efect of soil properties on vine performance’.

van Leeuwen, C. (2010) ‘9 - Terroir: the efect of the physical environment on vine growth, grape ripening and wine sensory attributes’, in A.G. Reynolds (ed.) Managing Wine Quality. Woodhead Publishing (Woodhead Publishing Series in Food Science, Technology and Nutrition), pp. 273–315. Available at: 10.1533/9781845699284.3.273.

Li, Y.-B. et al. (2022) ‘Appropriate Soil Heat Treatment Promotes Growth and Disease Suppression of Panax notoginseng by Interfering with the Bacterial Community’, Journal of Microbiology and Biotechnology, 32(3), pp. 294–301. Available at: 10.4014/jmb.2112.12005.

Louca, S., Parfrey, L.W. and Doebeli, M. (2016) ‘Decoupling function and taxonomy in the global ocean microbiome’, Science (New York, N.Y.), 353(6305), pp. 1272–1277. Available at: 10.1126/science.aaf4507.

Love, M.I., Huber, W. and Anders, S. (2014) ‘Moderated estimation of fold change and dispersion for RNA-seq data with DESeq2’, Genome Biology, 15(12), p. 550. Available at: 10.1186/s13059-014-0550-8.

Marasco, R. et al. (2013) ‘Plant growth promotion potential is equally represented in diverse grapevine root-associated bacterial communities from diferent biopedoclimatic environments’, BioMed Research International, 2013, p. 491091. Available at: 10.1155/2013/491091.

Marasco, R. et al. (2022) ‘Rootstock–scion combination contributes to shape diversity and composition of microbial communities associated with grapevine root system’, Environmental Microbiology, 24(8), pp. 3791–3808. Available at: 10.1111/1462-2920.16042.

Mesny, F., Hacquard, S. and Thomma, B.P. (2023) ‘Co-evolution within the plant holobiont drives host performance’, EMBO reports, 24(9), p. e57455. Available at: 10.15252/embr.202357455.

Nerva, L. et al. (2021) ‘Microscale analysis of soil characteristics and microbiomes reveals potential impacts on plants and fruit: vineyard as a model case study’, Plant and Soil, 462(1), pp. 525–541. Available at: 10.1007/s11104-021-04884-2.

Nguyen, N.H. et al. (2016) ‘FUNGuild: An open annotation tool for parsing fungal community datasets by ecological guild’, Fungal Ecology, 20, pp. 241–248. Available at: 10.1016/j.funeco.2015.06.006.

Oksanen, J. et al. (2024) ‘vegan: Community Ecology Package’. Available at: https://cran.r-project.org/web/packages/vegan/index.html (Accessed: 25 July 2024).

Pascale, A. et al. (2020) ‘Modulation of the Root Microbiome by Plant Molecules: The Basis for Targeted Disease Suppression and Plant Growth Promotion’, Frontiers in Plant Science, 10, p. 1741. Available at: 10.3389/fpls.2019.01741.

Quast, C. et al. (2013) ‘The SILVA ribosomal RNA gene database project: improved data processing and web-based tools’, Nucleic Acids Research, 41(D1), pp. D590–D596. Available at: 10.1093/nar/gks1219.

Quezada, C. et al. (2014) ‘Influence of Soil Physical Properties on Grapevine Yield and Maturity Components in an Ultic Palexeralf Soils, Central-Southern, Chile’, Open Journal of Soil Science, 4(4), pp. 127–135. Available at: 10.4236/ojss.2014.44016.

R Core Team (2021) R: A Language and Environment for Statistical Computing. Vienna, Austria: R Foundation for Statistical Computing. Available at: https://www.R-project.org/.

Redmile-Gordon, M., White, R.P. and Brookes, P.C. (2011) ‘Evaluation of substitutes for paraquat in soil microbial ATP determinations using the trichloroacetic acid based reagent of Jenkinson and Oades (1979)’, Soil Biology and Biochemistry, 43(5), pp. 1098– 1100. Available at: 10.1016/j.soilbio.2011.01.007.

Rhoades, J.D. (1996) ‘Salinity: Electrical Conductivity and Total Dissolved Solids’, in Methods of Soil Analysis. John Wiley & Sons, Ltd, pp. 417–435. Available at: 10.2136/sssabookser5.3.c14.

Rolli, E. et al. (2017) ‘Root-associated bacteria promote grapevine growth: from the laboratory to the field’, Plant and Soil, 410(1), pp. 369–382. Available at: 10.1007/s11104-016-3019-6.

Sánchez-Cañizares, C. et al. (2017) ‘Understanding the holobiont: the interdependence of plants and their microbiome’, Current Opinion in Microbiology, 38, pp. 188–196. Available at: 10.1016/j.mib.2017.07.001.

Turner, T.R., James, E.K. and Poole, P.S. (2013) ‘The plant microbiome’, Genome Biology, 14(6), p. 209. Available at: 10.1186/gb-2013-14-6-209.

Uyan, M., Janus, J. and Ertunç, E. (2023) ‘Land Use Suitability Model for Grapevine (Vitis vinifera L.) Cultivation Using the Best Worst Method: A Case Study from Ankara/Türkiye’, Agriculture, 13(9), p. 1722. Available at: 10.3390/agriculture13091722.

Vandenkoornhuyse, P. et al. (2015) ‘The importance of the microbiome of the plant holobiont’, New Phytologist, 206(4), pp. 1196–1206. Available at: 10.1111/nph.13312.

Visconti, F., López, R. and Olego, M.Á. (2024) ‘The Health of Vineyard Soils: Towards a Sustainable Viticulture’, Horticulturae, 10(2), p. 154. Available at: 10.3390/horticulturae10020154.

Vukicevich, E. et al. (2018) ‘Groundcover management changes grapevine root fungal communities and plant-soil feedback’, Plant and Soil, 424(1), pp. 419–433. Available at: 10.1007/s11104-017-3532-2.

Waite, H. and May, P. (2005) ‘The efects of hot water treatment, hydration and order of nursery operations on cuttings of Vitis vinifera cultivars’, Phytopathologia Mediterranea, 44(2), pp. 144–152.

Wang, R., Sun, Q. and Chang, Q. (2015) ‘Soil Types Efect on Grape and Wine Composition in Helan Mountain Area of Ningxia’, PLOS ONE, 10(2), p. e0116690. Available at: 10.1371/journal.pone.0116690.

Wang, W. et al. (2025) ‘Soil organic matter composition afects ecosystem multifunctionality by mediating the composition of microbial communities in long-term restored meadows’, Environmental Microbiome, 20(1), p. 22. Available at: 10.1186/s40793-025-00678-6.

Wankhade, A. et al. (2025) ‘A Review of Plant–Microbe Interactions in the Rhizosphere and the Role of Root Exudates in Microbiome Engineering’, Applied Sciences, 15(13), p. 7127. Available at: 10.3390/app15137127.

Wei, T. et al. (2017) ‘Package “corrplot”’, Statistician, 56(316), p. e24.

Wilkinson, L. (2011) ‘ggplot2: Elegant Graphics for Data Analysis by WICKHAM, H.’, Biometrics, 67(2), pp. 678–679. Available at: 10.1111/j.1541-0420.2011.01616.x.

Zarraonaindia, I. et al. (2015) ‘The Soil Microbiome Influences Grapevine-Associated Microbiota’, mBio, 6(2), pp. e02527–14. Available at: 10.1128/mBio.02527-14.

