## Supplementary material for "Variations in the root-soil system influence the grapevine holobiont by shaping plant physiology and root microbiome"

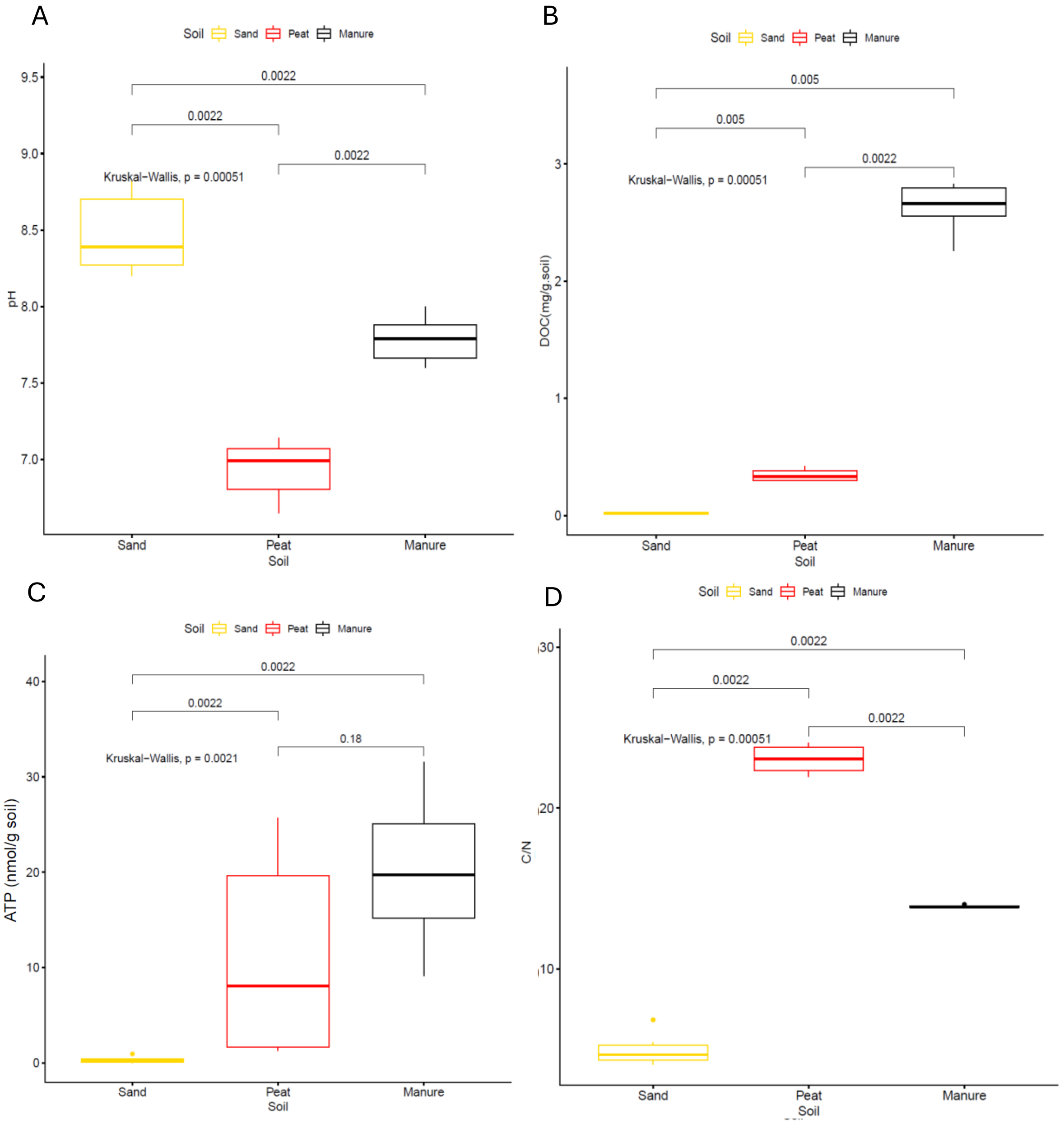


Supplementary Fig. S1. Boxplot with pairwise-Wilcoxon test (in brackets) showing the physico-chemical parameters of the three soils at T0. (A) pH level, (B) Dissolved organic carbon level, (C) Adenosine Triphosphate

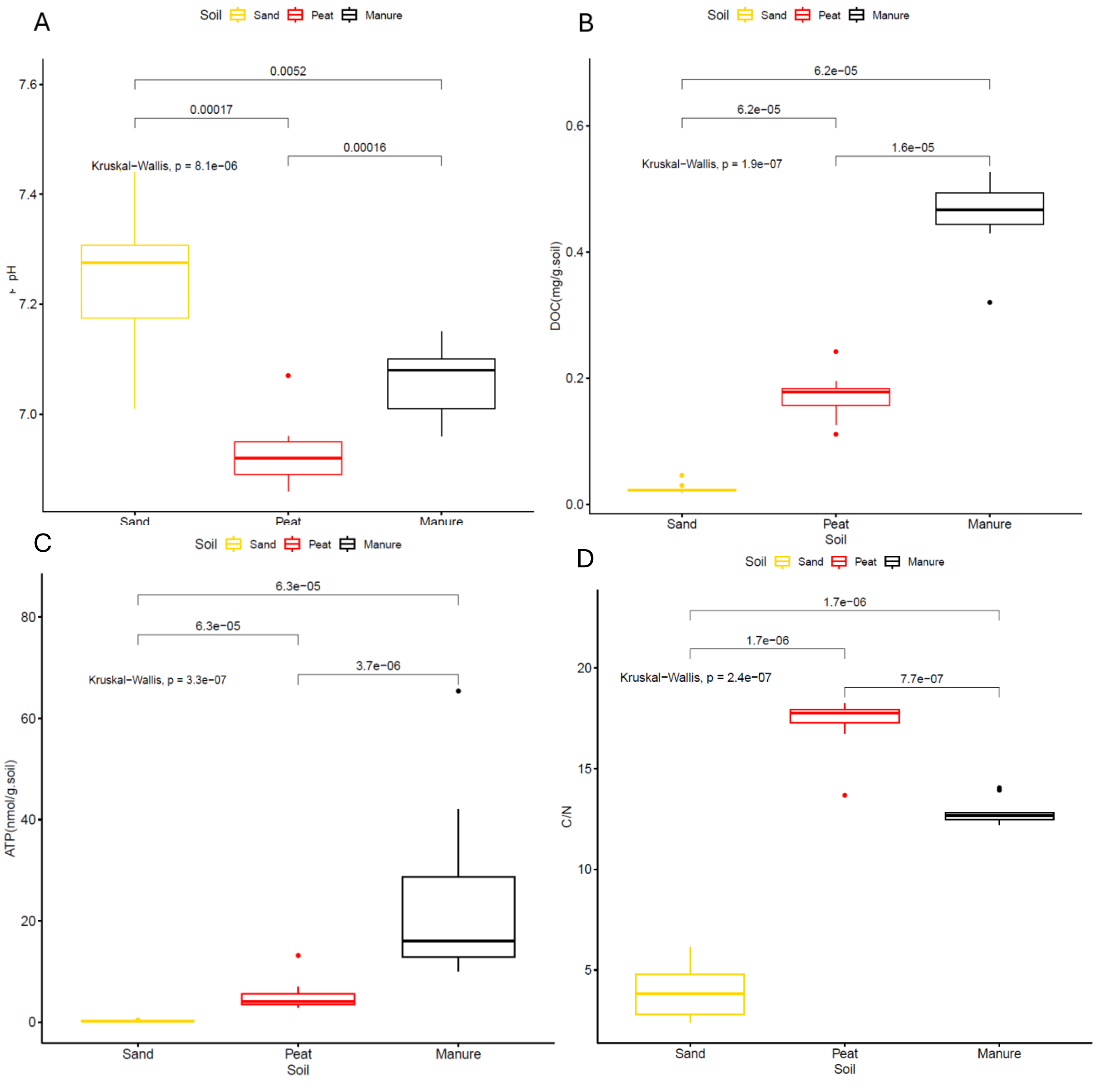


Supplementary Fig. S2: Boxplot with pairwise-Wilcoxon test (in brackets) showing the physico-chemical parameters of the three soil types at T1. (A) pH, (B) Dissolved organic carbon (DOC) concentration, (C) Adenosine Triphosphate (ATP) concentration, (D) Organic carbon/organic nitrogen (C/N) ratio.

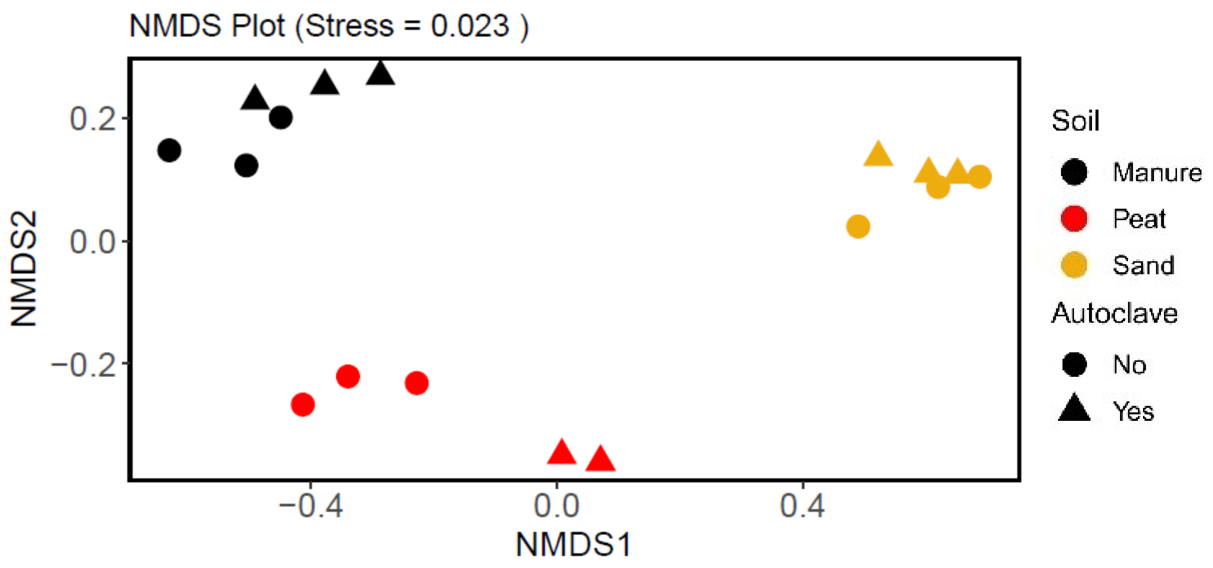


Supplementary Fig. S3: NMDS analysis based on physico-chemical parameters at T0.

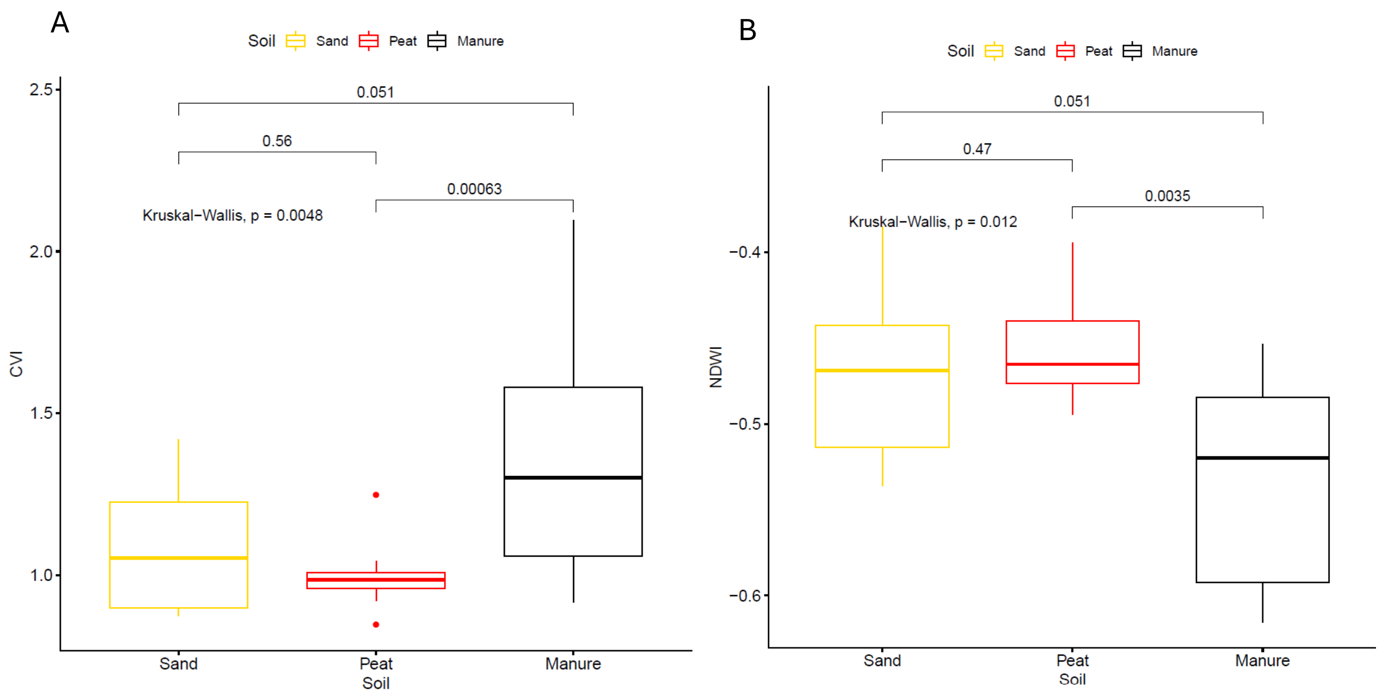


Supplementary Fig. S4: Boxplot with pairwise-Wilcoxon test (in brackets) showing the multispectral indices levels of the plants at T1 which present significant differences. (A) CVI, (B) NDWI.

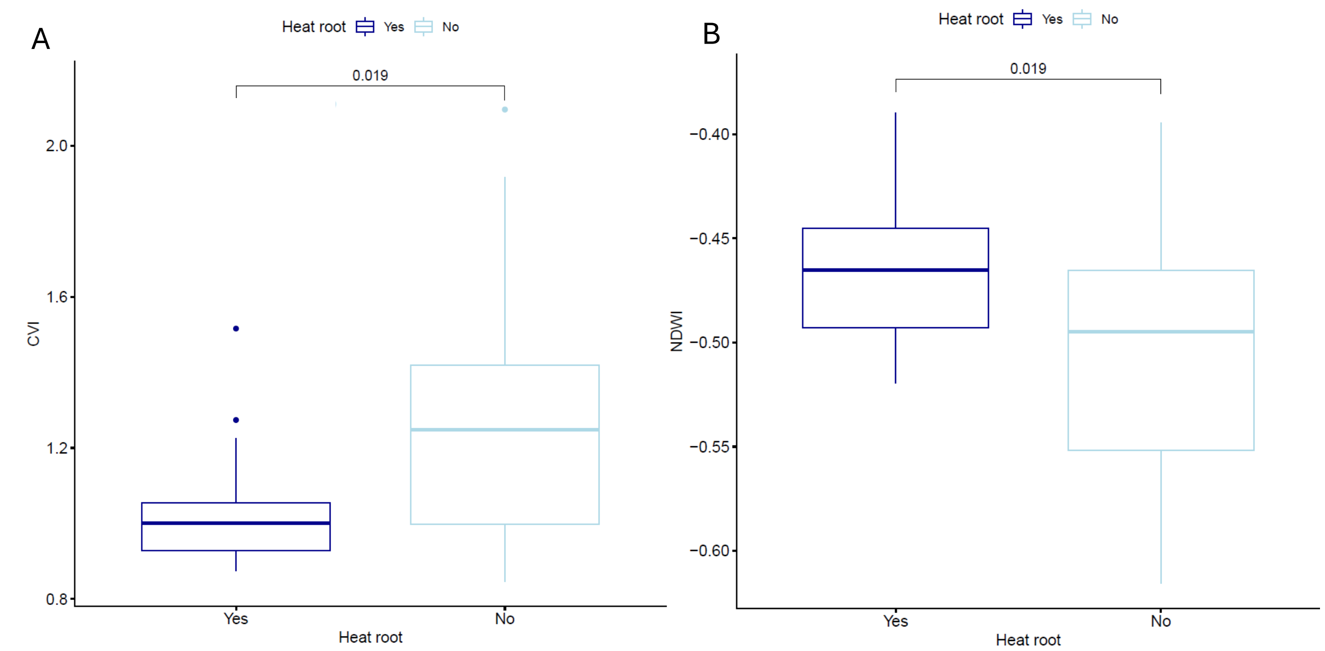


Supplementary Fig. S5: Boxplot with pairwise-Wilcoxon test (in brackets) showing the multispectral indices levels of the plants at T1 in the heat root/not heat root treatment which present significant differences. (A) CVI, (B) NDWI.

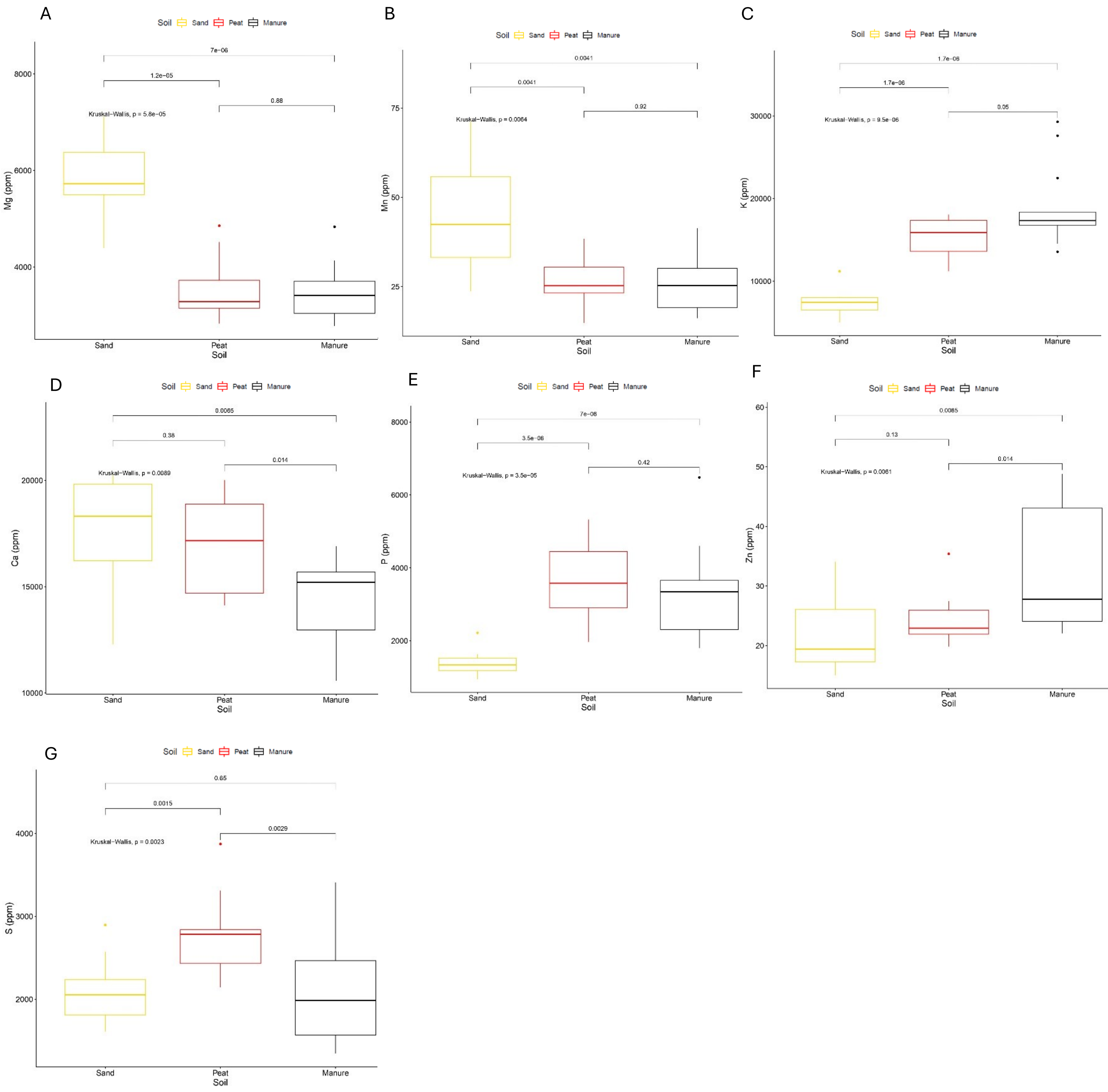


Supplementary Fig. S6: Boxplot with pairwise-Wilcoxon test (in brackets) showing the multielemental concentration, measured in mg/kg, in the leaves of the plants at T1 in the three types of soil which present significant differences. (A) magnesium (Mg), (B) manganese (Mn), (C) potassium (K), (D) calcium (Ca),(E) phosphorus, (F) zinc (Zn), (G) sulphur (S) .

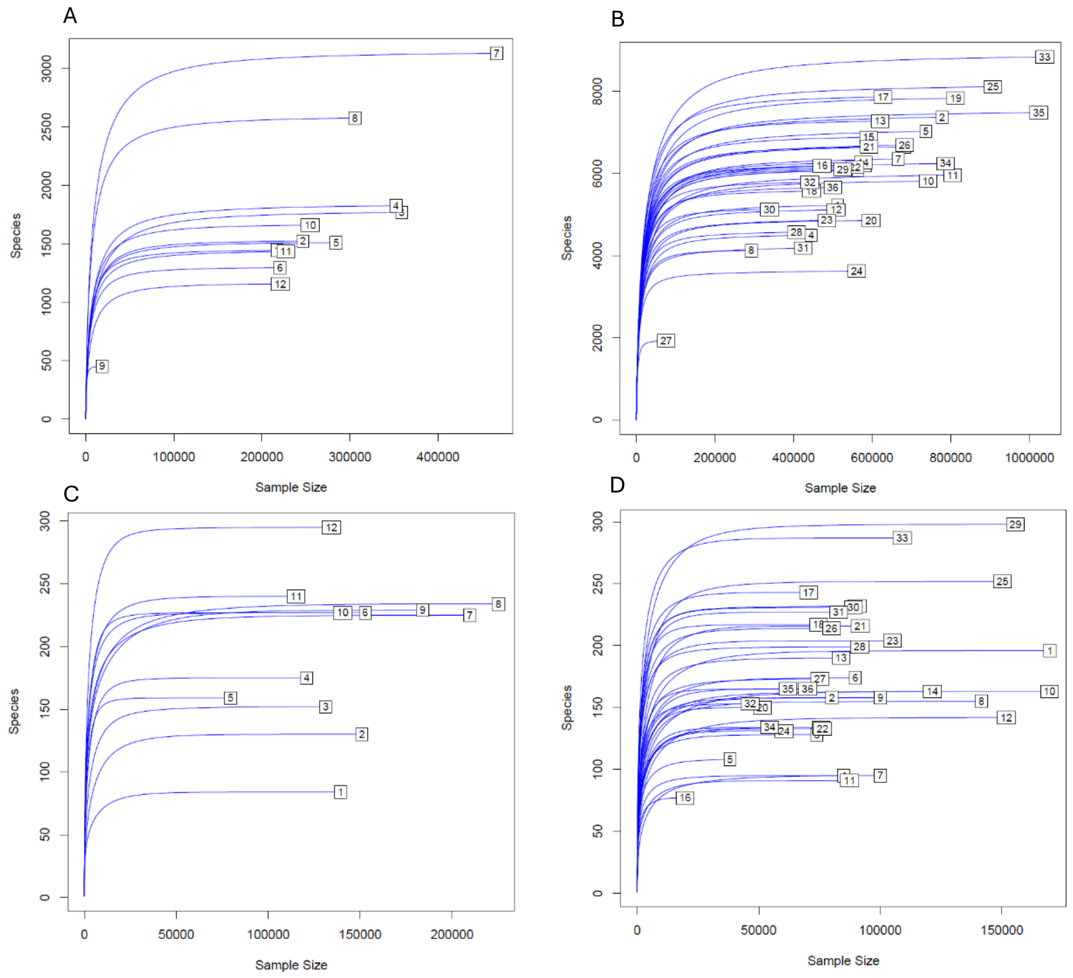


Supplementary Fig. S7: Rarefaction curves of the T0 16s sequencing (A), T1 16s sequencing (B), T0 ITS sequencing (C), T0 ITS sequencing (D).

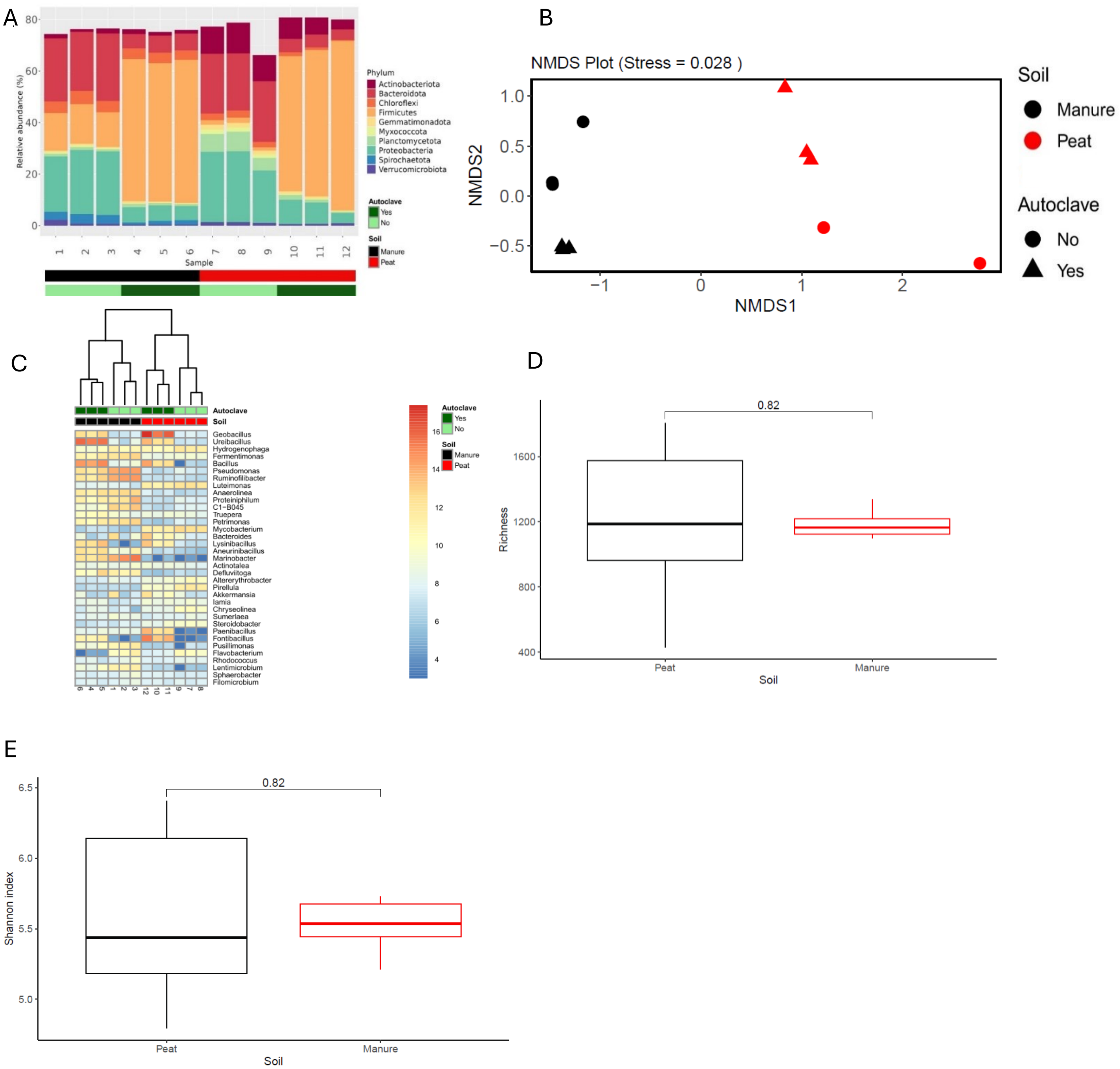


Supplementary Fig. S8: Soil bacterial microbiome at T0. (A) Relative abundance of the most predominant phyla. (B) NMDS analysis of the bulk soil bacterial microbiome. (C) Clustering and representation of the most abundant genera. Boxplot with pairwise- Wilcoxon test (in brackets) of the (D) Richness and (E) Shannon index in the two types of soil.

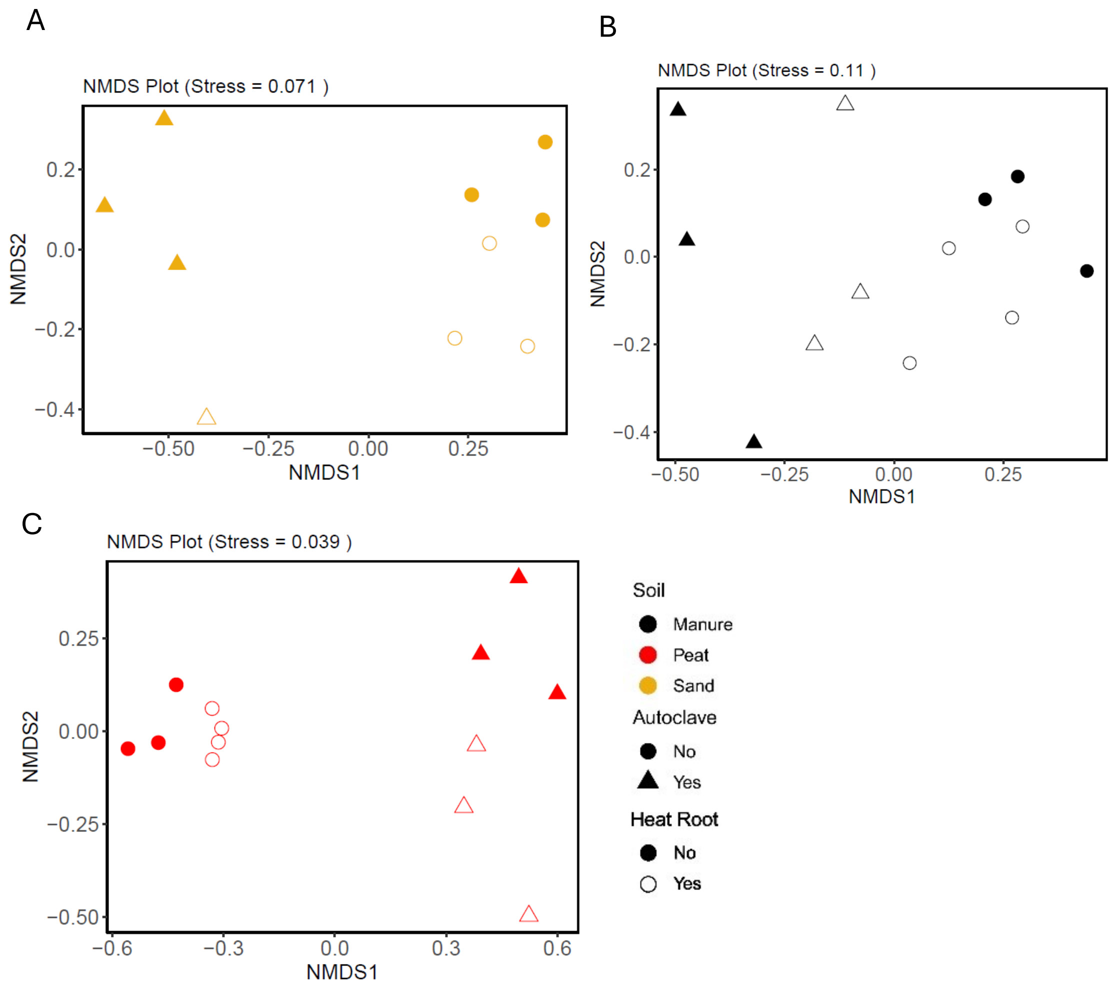


Supplementary Fig. S9: NMDS plot of the T1 bacterial microbiome (A) Sand, (B) Manure, (C) Peat.

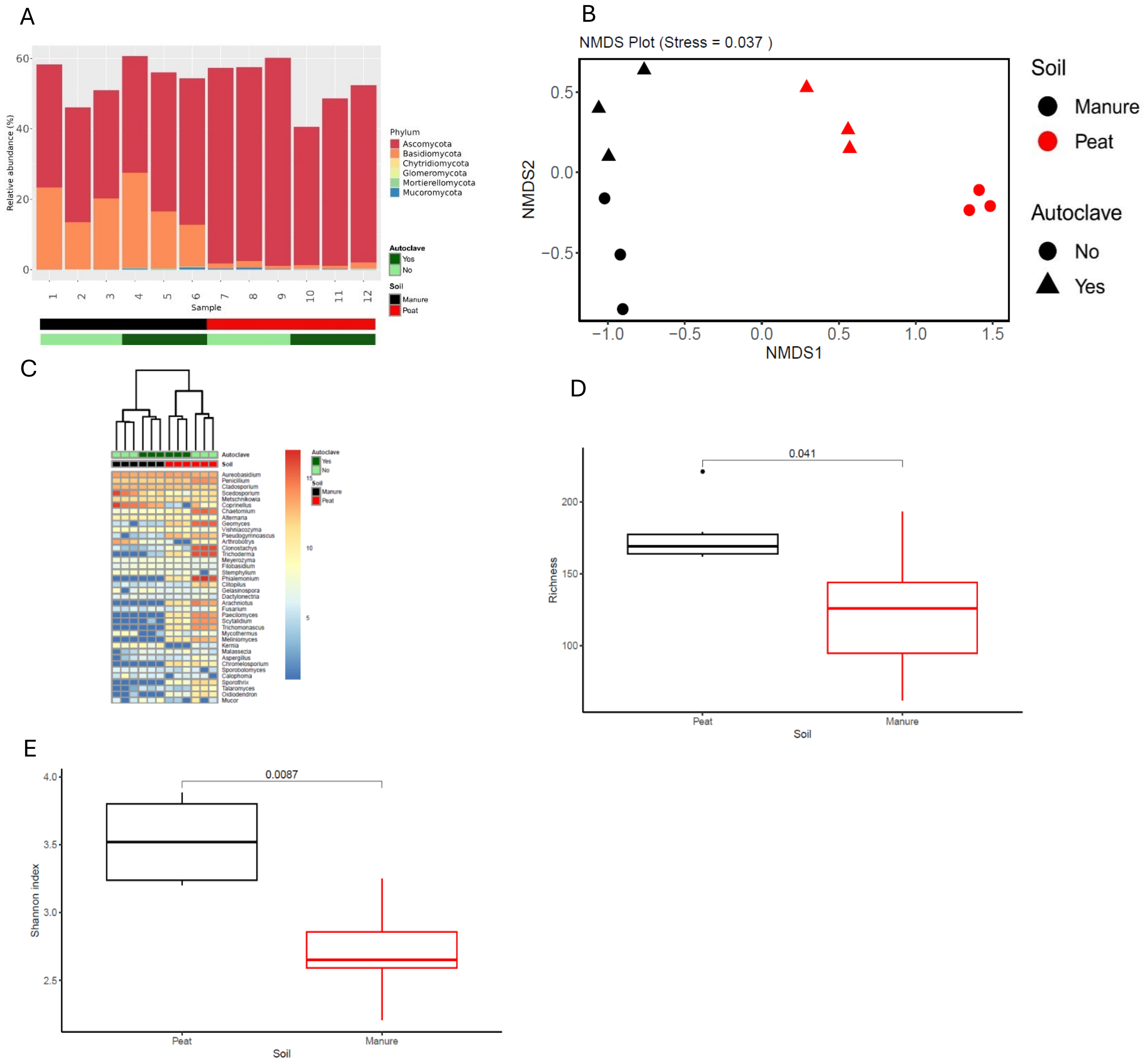


Supplementary Fig. S10: Soil fungal microbiome at T0. (A) Relative abundance of the most predominant phyla. (B) Heatmap of the 35 most abundant genera with clustering analysis. (B) NMDS analysis of the bulk soil fungal microbiome. Boxplot with pairwise- Wilcoxon test (in brackets) of the (D) Richness and (E) Shannon index in the two types of soil.

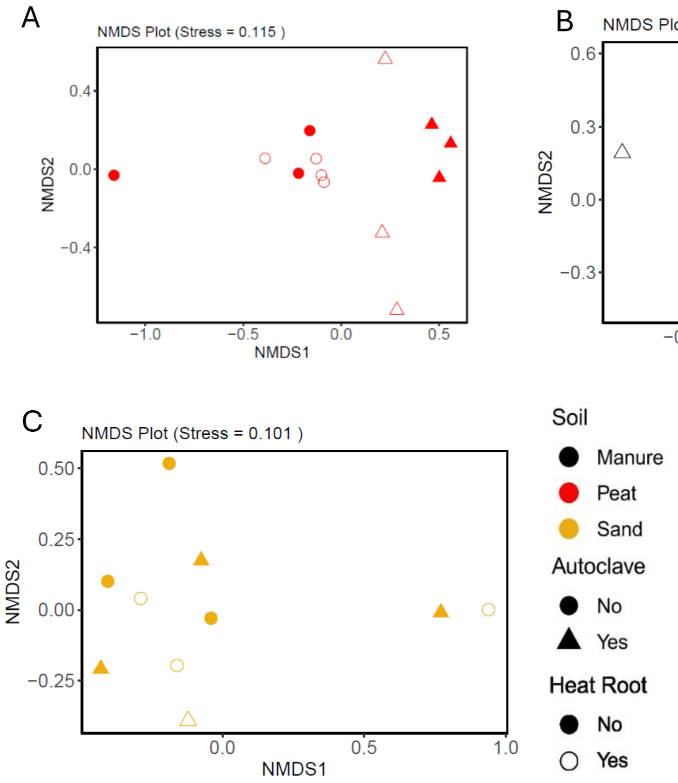


Supplementary Fig. S11: NMDS plot of the T1 fungal microbiome (A) Sand, (B) Manure, (C) Peat.

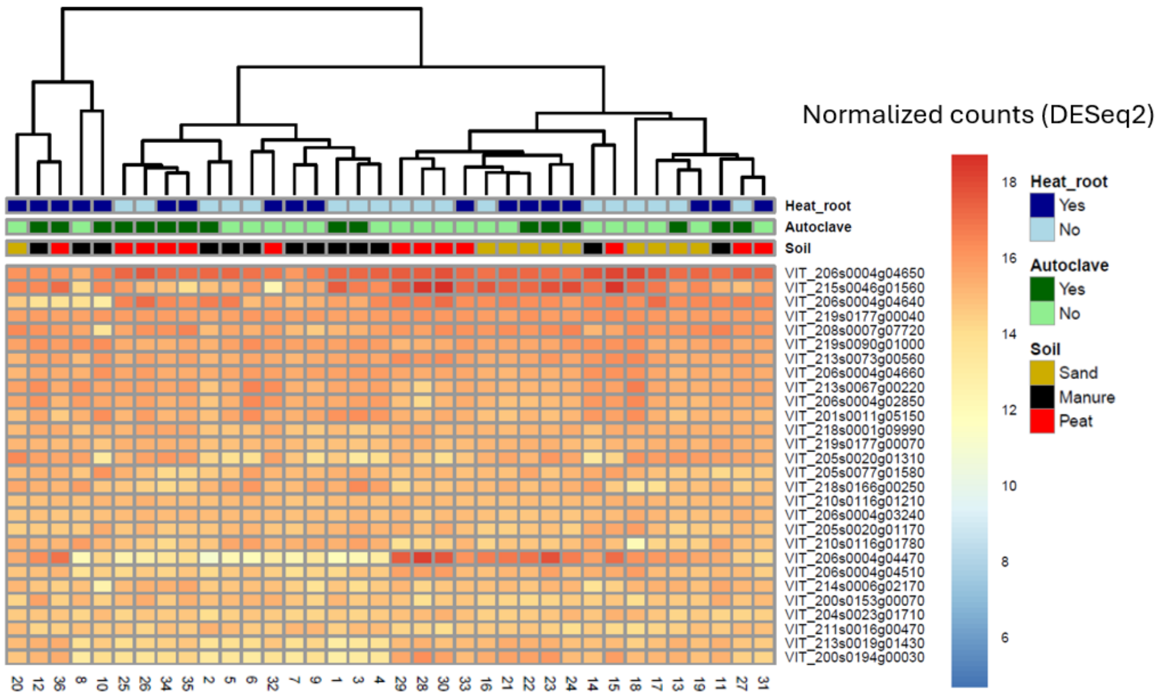


Supplementary Fig. S12: Heatmap and clustering of the normalized counts of the 25 most expressed genera in the roots. Clustering is performed considering the all the expressed genes.

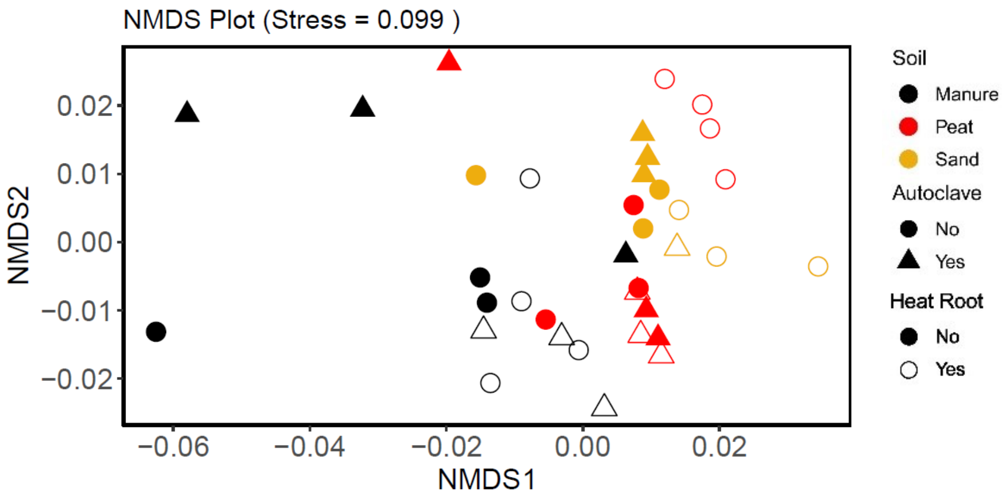


Supplementary Fig. S13: NMDS plot performed transcriptomic data.

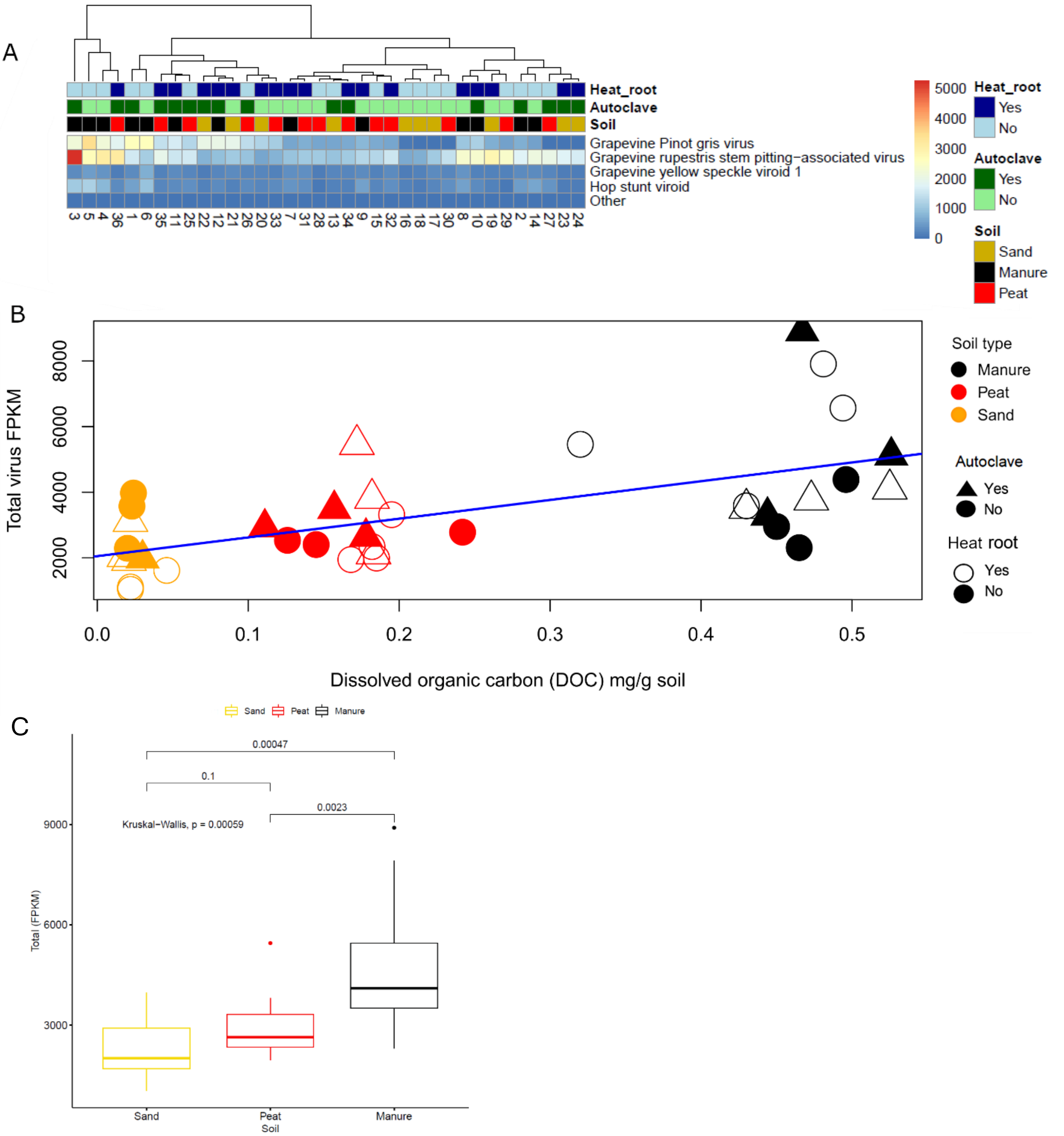


Supplementary Fig. S14: (A) Heatmap showing the relative abundance of the identified virus expressed in Fragment per Kilobase Million with clustering analysis per type of soil. (B) Total viral abundance in root as a function of DOC concentration in soil) (C) Boxplot with Wilcoxon test (in brackets) comparing total viral abundance in the three types of soil.

Supplementary Table S1: Categories considered for functional analysis. FAPROTAX categories: Categories considered for bacterial functional analysis. FUNGuild categories: categories considered for fungal functional analysis.

| FAPROTAX categories | FUNGuild categories |
|---|---|
| Aerobic Ammonia Oxidation, Aerobic Nitrite Oxidation, Nitrification, Nitrate Denitrification, Nitrite Denitrification, Nitrous Oxide Denitrification, Denitrification, Nitrite Respiration, Nitrogen Fixation, Xylanolysis, Fermentation, Plant Pathogen, Nitrate Respiration, Nitrogen Respiration, Nitrate Reduction, Ureolysis. | Arbuscular Mycorrhizal, Undefined Saprotroph, Wood Saprotroph, Plant Pathogen, Plant Saprotroph, Dung Saprotroph, Epiphyte, Pollen Saprotroph, Orchid Mycorrhizal, Lichen Parasite, Fungal Parasite, Ectomycorrhizal, and Ericoid Mycorrhizal. |

Supplementary Table S2: Results of Pearson correlation tests between the soil physico-ochemical properties at T0. The lower diagonal contains the Pearson correlation coefficient (r), the upper diagonal contains the p-values. **ATP**: adenosine triphosphate content, **pH**: soil pH, **EC**: electrical conductivity, **DOC**: dissolved organic carbon, **DN**: dissolved nitrogen, **TOC**: total organic carbon, **TN**: total nitrogen, **C/N ratio**: carbon-to-nitrogen ratio.

| | ATP | pH | EC | DOC | DN | TOC | TN | C/N ratio |
|---|---|---|---|---|---|---|---|---|
| ATP | | 0.135 | 0.004 | 0.004 | 0.0005 | 0.0006 | 0.0001 | 0.077 |
| pH | -0.366 | | 0.555 | 0.827 | 0.173 | 0.634 | 0.284 | 1.060E-09 |
| EC | 0.632 | -0.149 | | 7.690E-10 | 3.808E-08 | 1.143E-08 | 3.967E-08 | 0.485 |
| DOC | 0.632 | -0.055 | 0.955 | | 2.443E-09 | 7.523E-13 | 6.568E-10 | 0.687 |
| DN | 0.734 | -0.336 | 0.925 | 0.947 | | 1.206E-09 | 4.484E-12 | 0.103 |
| TOC | 0.726 | -0.120 | 0.936 | 0.981 | 0.952 | | 3.375E-14 | 0.496 |
| TN | 0.778 | -0.267 | 0.925 | 0.956 | 0.976 | 0.987 | | 0.189 |
| C/N ratio | 0.427 | -0.953 | 0.176 | 0.102 | 0.397 | 0.171 | 0.324 | |

Supplementary Table S3: Pearson correlation tests between the soil physiochemical properties at T1. The lower diagonal contains the Pearson correlation coefficient (r), the upper diagonal contains the p-values. **ATP**: adenosine triphosphate content, **pH**: soil pH, **EC**: electrical conductivity, **DOC**: dissolved organic carbon, **DN**: dissolved nitrogen, **TOC**: total organic carbon, **TN**: total nitrogen, **C/N ratio**: carbon-to-nitrogen ratio.

| | pH | EC | DOC | DN | TN | TOC | C/N ration | ATP |
|---|---|---|---|---|---|---|---|---|
| pH | | 0.428 | 0.082 | 0.250 | 0.433 | 0.230 | 7.137E-09 | 0.636 |
| EC | -0.136 | | 0 | 0 | 0 | 0 | 0.182 | 1.089E-07 |
| DOC | -0.294 | 0.961 | | 0 | 0 | 0.010 | 0.010 | 0.686 |
| DN | -0.197 | 0.948 | 0.962 | | 0 | 0.059 | 0.059 | 0.630 |
| TN | -0.135 | 0.991 | 0.964 | 0.939 | | 0.128 | 0.128 | 0.770 |
| TOC | -0.205 | 0.984 | 0.975 | 0.947 | 0.994 | | 0.037 | 0.764 |
| C/N | -0.796 | 0.228 | 0.423 | 0.318 | 0.258 | 0.348 | | 0.215 |
| ATP | -0.081 | 0.754 | 0.686 | 0.630 | 0.770 | 0.764 | 0.215 | |

Supplementary Table S4: Nested PERMANOVA of soil p**hysico-chemical** parameters (DOC, pH, ATP, and C/N) at T0 (formula = diss_matrix ~ Soil/Autoclave). **Soil**: Effect of soil type. **Soil:Autoclave**: Nested effect of autoclaving within each soil type. **Df**: Degrees of freedom. **SumOfSqs**: Sum of squares. **$R^2$**: Proportion of variance explained. **F**: F-statistic. **Pr(>F)**: p-value.

| | Df | SumOfSqs | R2 | F | Pr(>F) |
|---|---|---|---|---|---|
| Soil | 2 | 1.291 | 0.890 | 177.939 | 0.001 |
| Soil:Autoclave | 3 | 0.115 | 0.080 | 10.608 | 0.001 |
| Residual | 12 | 0.043 | 0.030 | | |
| Total | 17 | 1.450 | 1.000 | | |

Supplementary Table S5: Nested PERMANOVA of soil p**hysico-chemical** parameters (DOC, pH, ATP, and C/N) at T1 (formula = diss_matrix ~ Soil/Autoclave/Heat root). **Soil**: Effect of soil type **Soil:Autoclave**: Nested effect of autoclaving within each soil type. **Soil:Autoclave:Heat Root**: Nested effect of root heat treatment within each soil-autoclave combination. **Df**: Degrees of freedom. **SumOfSqs**: Sum of squares. **$R^2$**: Proportion of variance explained. **F**: F-statistic. **Pr(>F)**: p-value.

| | Df | SumOfSqs | R2 | F | Pr(>F) |
|---|---|---|---|---|---|
| Soil | 2 | 2.250 | 0.863 | 144.414 | 0.001 |
| Soil:Autoclave | 3 | 0.082 | 0.031 | 3.493 | 0.015 |
| Soil:Autoclave:Heat Root | 6 | 0.089 | 0.034 | 1.913 | 0.065 |
| Residual | 24 | 0.187 | 0.071 | | |
| Total | 35 | 2.608 | 1.000 | | |

Supplementary table S6: Nested PERMANOVA of the multispectral indices NDVI, CVI, GDVI, NDWI at T1 (formula = diss_matrix ~ Soil/Autoclave/Heat root). **Soil**: Effect of soil type. **Soil:Autoclave**: Nested effect of autoclaving within each soil type. **Soil:Autoclave:Heat Root**: Nested effect of root heat treatment within each soil-autoclave combination. **Df**: Degrees of freedom. **SumOfSqs**: Sum of squares. **R²**: Proportion of variance explained. **F**: F-statistic. **Pr(>F)**: p-value.

| | Df | SumOfSqs | R2 | F | Pr(>F) |
|---|---|---|---|---|---|
| Soil | 2 | 0.185 | 0.274 | 8.504 | 0.001 |
| Soil:Autoclave | 3 | 0.073 | 0.109 | 2.257 | 0.079 |
| Soil:Autoclave:Heat Root | 5 | 0.155 | 0.229 | 2.851 | 0.02 |
| Residual | 24 | 0.261 | 0.3868 | | |
| Total | 34 | 0.673 | 1 | | |

Supplementary table S7: Nested PERMANOVA of the multielemental ions Ca, Cu, Fe, K, Mg, Mn, Na, P, S, Zn at T1 (formula = diss_matrix ~ Soil/Autoclave/Heat root). **Soil**: Effect of soil type. **Soil:Autoclave**: Nested effect of autoclaving within each soil type. **Soil:Autoclave:Heat Root**: Nested effect of root heat treatment within each soil-autoclave combination. **Df**: Degrees of freedom. **SumOfSqs**: Sum of squares. **R²**: Proportion of variance explained. **F**: F-statistic. **Pr(>F)**: p-value.

| | Df | SumOfSqs | R2 | F | Pr(>F) |
|---|---|---|---|---|---|
| Soil | 2 | 0.380 | 0.623 | 30.208 | 0.001 |
| Soil:Autoclave | 3 | 0.032 | 0.053 | 1.712 | 0.143 |
| Soil:Autoclave:Heat Root | 6 | 0.046 | 0.076 | 1.226 | 0.311 |
| Residual | 24 | 0.151 | 0.248 | | |
| Total | 35 | 0.610 | 1.000 | | |

Supplementary Table S8: Summary of sequencing data for 16S and ITS amplicons across time points. **Amplicon Type**: Targeted marker (16S for bacteria, ITS for fungi). **Time Point**: Sampling time (T0 = initial, T1 = final). **Total Reads**: Total number of sequences obtained for each amplicon type and time point. **Min Reads per Sample**: Lowest number of reads obtained in an individual sample. **Max Reads per Sample**: Highest number of reads obtained in an individual sample. **Average Reads per Sample**: Mean number of reads per sample. **Amplicon Sequence Variants**: Number of unique ASVs detected.

| Amplicon Type | Time Point | Total Reads | Min Reads per Sample | Max Reads per Sample | Average Reads per Sample | Amplicon Sequence Variants |
|---|---|---|---|---|---|---|
| 16S | T0 | 3,794,673 | 27,120 | 547,314 | 316,223 | 3,533 |
| 16S | T1 | 24,597,687 | 117,169 | 1,162,043 | 683,269 | 11,308 |
| ITS | T0 | 2,605,034 | 126,286 | 298,214 | 217,086 | 600 |
| ITS | T1 | 8,482,398 | 9,754 | 442,723 | 235,622 | 774 |

Supplementary table S9: Nested PERMANOVA of the ASVs of the T0 rhizosphere bacterial microbiome (formula = diss_matrix ~ Soil / Autoclave). **Soil**: Effect of soil type. **Soil:Autoclave**: Nested effect of autoclaving within each soil type. **Df**: Degrees of freedom. **SumOfSqs**: Sum of squares. **R²**: Proportion of variance explained. **F**: F-statistic. **Pr(>F)**: p-value.

| | Df | SumOfSqs | R2 | F | Pr(>F) |
|---|---|---|---|---|---|
| Soil | 1 | 1.643 | 0.490 | 58.994 | 0.001 |
| Soil:Autoclave | 2 | 1.489 | 0.444 | 26.739 | 0.001 |
| Residual | 8 | 0.223 | 0.066 | | |
| Total | 11 | 3.356 | 1 | | |

Supplementary table S10: Nested PERMANOVA of the ASVs table of the T1 rhizosphere bacterial microbiome (formula = diss_matrix ~ Soil / Autoclave,/ Heat root). **Soil**: Effect of soil type. **Soil:Autoclave**: Nested effect of autoclaving within each soil type. **Soil:Autoclave:Heat Root**: Nested effect of root heat treatment within each soil-autoclave combination. **Df**: Degrees of freedom. **SumOfSqs**: Sum of squares. **R²**: Proportion of variance explained. **F**: F-statistic. **Pr(>F)**: p-value.

| | Df | SumOfSqs | R2 | F | Pr(>F) |
|---|---|---|---|---|---|
| Soil | 2 | 4.759 | 0.427 | 17.417 | 0.001 |
| Soil:Autoclave | 3 | 1.760 | 0.158 | 4.293 | 0.001 |
| Soil:Autoclave:Heat Root | 6 | 1.337 | 0.120 | 1.631 | 0.005 |
| Residual | 24 | 3.279 | 0.294 | | |
| Total | 35 | 11.134 | 1.000 | | |

Supplementary table S11: Nested PERMANOVA of ASV of the T0 bulk soil fungal microbiome (formula = diss_matrix ~ Soil / Autoclave). **Soil**: Effect of soil type. **Soil:Autoclave**: Nested effect of autoclaving within each soil type. **Soil:Autoclave:Heat Root**: Nested effect of root heat treatment within each soil-autoclave combination. **Df**: Degrees of freedom. **SumOfSqs**: Sum of squares. **R²**: Proportion of variance explained. **F**: F-statistic. **Pr(>F)**: p-value.

| | Df | SumOfSqs | R2 | F | Pr(>F) |
|---|---|---|---|---|---|
| Soil | 1 | 1.506 | 0.573 | 48.143 | 0.001 |
| Soil:Autoclave | 2 | 0.872 | 0.332 | 13.942 | 0.002 |
| Residual | 8 | 0.250 | 0.095 | | |
| Total | 11 | 2.629 | 1.000 | | |

Supplementary table S12: Nested PERMANOVA of ASV of the T1 rhizosphere fungal microbiome (formula = diss_matrix ~ Soil / Autoclave / Heat root). **Soil**: Effect of soil type. **Soil:Autoclave**: Nested effect of autoclaving within each soil type.

**Soil:Autoclave:Heat Root**: Nested effect of root heat treatment within each soil-autoclave combination. **Df**: Degrees of freedom. **SumOfSqs**: Sum of squares. **R²**: Proportion of variance explained. **F**: F-statistic. **Pr(>F)**: p-value.

| | Df | SumOfSqs | $R^2$ | F | Pr(>F) |
|---|---|---|---|---|---|
| Soil | 2 | 2.118 | 0.185 | 4.559 | 0.001 |
| Soil:Autoclave | 3 | 1.471 | 0.128 | 2.111 | 0.001 |
| Soil:Autoclave:Heat Root | 6 | 2.299 | 0.201 | 1.650 | 0.002 |
| Residual | 24 | 5.574 | 0.486 | | |
| Total | 35 | 11.462 | 1.000 | | |

Supplementary table S13: Nested PERMANOVA of the root transcriptome at T1 (formula = diss_matrix ~ Soil / Autoclave / Heat root). **Soil**: Effect of soil type. **Soil:Autoclave**: Nested effect of autoclaving within each soil type. **Soil:Autoclave:Heat Root**: Nested effect of root heat treatment within each soil-autoclave combination. **Df**: Degrees of freedom. **SumOfSqs**: Sum of squares. **R²**: Proportion of variance explained. **F**: F-statistic. **Pr(>F)**: p-value.

| | Df | SumOfSqs | R2 | F | Pr(>F) |
|---|---|---|---|---|---|
| Soil | 2 | 0.129 | 0.227 | 6.76 | 0.001 |
| Soil:Autoclave | 3 | 0.064 | 0.113 | 2.243 | 0.011 |
| Soil:Autoclave:Heat Root | 6 | 0.146 | 0.257 | 2.55 | 0.002 |
| Residual | 24 | 0.229 | 0.403 | | |
| Total | 35 | 0.567 | 1 | | |
